# p53 engages the cGAS/STING cytosolic DNA sensing pathway for tumor suppression

**DOI:** 10.1101/2022.05.26.493595

**Authors:** Monisankar Ghosh, Suchandrima Saha, Jinyu Li, David C Montrose, Luis A. Martinez

**Author notes:** Corresponding author: Luis A. Martinez.

## Abstract

Tumor suppression by p53 is known to involve cell autonomous and non-cell autonomous mechanisms. p53 has been shown to suppress tumor growth by modulating immune system functions, however, the mechanistic basis for this activity is not well understood. Here we report that p53 promotes the degradation of the DNA exonuclease TREX1, resulting in cytosolic dsDNA accumulation. We demonstrate that p53 requires the ubiquitin ligase TRIM24 to induce TREX1 ubiquitin-dependent degradation. The accumulation of cytosolic DNA due to p53’s suppression of TREX1 activates the cytosolic DNA sensor, cGAS and its downstream effectors STING/TBK1/IRF3 resulting in induction of Type I interferons. TREX1 overexpression sufficed to block WTp53 activation of the cGAS/STING pathway. WTp53 mediated induction of type I interferon (IFNB1) response could be suppressed by cGAS/STING knockout. We find that p53’s tumor suppressor activities are compromised by loss of signaling through the cGAS/STING pathway. Thus, our study reveals that p53 utilizes the cGAS/STING innate immune system pathway for both cell intrinsic and cell extrinsic tumor suppressor activities.

## Introduction

The product of the tumor suppressor gene, TP53, was initially identified as a cellular protein that interacts with viral proteins.^1–4^ Viral proteins often target cellular pathways that govern immune responses as a means of evading detection.^5–7^ WTp53 has been postulated to play a role in the control of the immune system, although the mechanism(s) and consequences of WTp53 induced crosstalk between tumor and immune cells is insufficiently understood.^8–14^ Given its multifaceted ability to suppress tumor growth and progression, the high frequency of TP53 mutations in human cancer contributes to the acquisition of several hallmarks of oncogenesis, among which is immune evasion.^15^ The cGAS/STING cytosolic DNA sensing pathway has emerged as a key mediator of the innate immune response.^16^ DNA is normally present in the nucleus and mitochondria, and the presence of cytosolic DNA is a danger-associated molecular pattern (DAMP) that is recognized by the pattern recognition receptor (PRR) cGAS.^17^ In homeostatic conditions, DNA exonucleases degrade cytoplasmic DNA to prevent the inappropriate activation of the cGAS/STING pathway.^18,19^ Cytoplasmic DNA is detected by cGAS, which becomes enzymatically active and produces a second messenger, cGAMP, which in turn triggers activation of downstream effectors of the pathway, STING/TBK1/IRF3.^20–22^ Importantly, epigenetic silencing of cGAS in cancer cells disables the innate immune response to the accumulation of cytosolic DNA.^23–26^ Loss of function in the cGAS pathway contributes to tumor development through cell-intrinsic and –extrinsic mechanisms.^26–30^ Mutant p53 has been shown to suppress downstream signaling from cGAS/STING pathway, thereby facilitating immune evasion.^31^ In contrast, wildtype p53 has been implicated in activating or contributing to downstream signaling from cGAS/STING although the mechanism remains unknown.^**31**,**32**^ Herein we report that WTp53 promotes the degradation of the DNA exonuclease TREX1, thereby resulting in the accumulation of cytoplasmic DNA which triggers activation of the cGAS/STING pathway. Notably, our data indicate that WTp53 employs the pathogen recognition receptor cGAS for tumor suppression.

## Results

### WTp53 activates the cGAS/STING innate immune pathway

We previously reported that mutant p53 can suppress downstream signaling from the cGAS/STING pathway.^31^ In contrast, we found that wildtype p53 activates the pathway.^31^ We extended this observation by examining whether wildtype p53 regulated downstream signaling in several different cell models in which we modulated p53 expression through different approaches. In three different mouse cell lines, we observed that wildtype p53 correlated with increased signaling of the cGAS/STING pathway as indicated by the increased phosphorylation of TBK1 substrates (phospho-TBK1, phospho-IRF3 and phospho-STING). Comparison of MEFs derived from p53 wildtype (p53^+/+^) or null (p53^−/−^) mice showed a clear positive correlation between WTp53 and cGAS/STING pathway. (Fig. 1A) We engineered p53 null 4T1 breast cancer cell line to inducibly express WTp53 and we observed that WTp53 expression increased TBK1 substrate phosphorylation. (Fig. 1B) Moreover, p53 knockdown with two different shRNAs reduced TBK1 substrate phosphorylation in CT26 murine colorectal cancer cells. (Fig. 1C) Our data indicates that WTp53 stimulates the cGAS/STING pathway.

**Figure 1:**
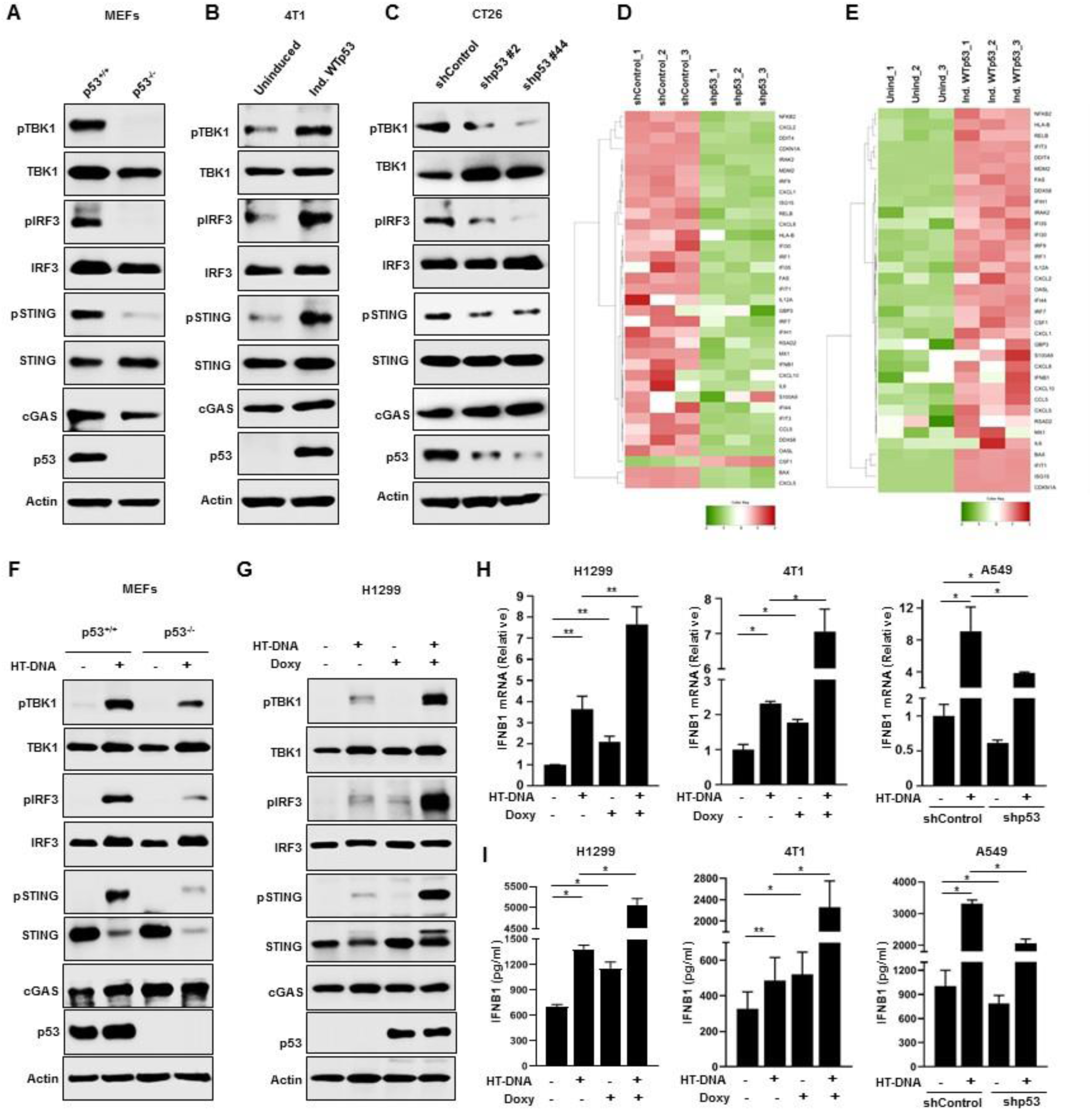
WTp53 activates the cGAS/STING innate immune pathway. Western blot analysis of (A) p53+/+ and p53-/- mouse embryonic fibroblast (MEFs). (B) P53 null mouse 4T1 cells engineered to inducibly express WTp53 were analyzed by western blot. (C) P53 was knocked down using two different shRNAs in CT26 cells and cell lysates were analyzed by western blot. (D-E) Heat maps show the differential gene expression as accessed by Nanostring (n=3) in (D) A549 shControl and shp53 and (E) H1299 Uninduced or induced WTp53. (F) p53+/+ and p53-/- MEFs were treated with 2 μg/ml of HT-DNA for 3 h and harvested for western blot analysis. (G) P53 null H1299 cells were engineered to inducibly express WT53. Cells were treated for 24 h with Doxycycline to induce p53 and then treated with 2 μg/ml of HT-DNA for 3 h and subjected to western blot analysis. (H and I) P53 null H1299 and 4T1 cells were induced to express WTp53, and A549 shControl or shp53 cells were treated with 2 μg/ml HT-DNA for 18 h, and cells were harvested for RT-PCR analysis of IFNB1 mRNA (H) or the conditioned medium was harvested for ELISA detection of secreted IFNB1 (I). Quantification graphs: In all panels, error bars represent mean with standard deviation. p values are based on Student’s t test. ***p < 0.001, **p < 0.01, *p < 0.05, ns=non-significant. See also Figure S1.

Since WTp53 activated the cGAS/STING pathway, we investigated if this resulted in differences in expression of interferon stimulated genes (ISG) via Nanostring analysis. We observed that WTp53 (in A549 and H1299 cells) had a high correlation with increased expression of ISGs. (Fig. 1D and 1E) We further validated the upregulation of key ISGs using real-time RT-PCR analysis of IFNB1, IFIT1, ISG15 and CXCL10 in H1299 induced WTp53, p53^+/+^ or p53^−/−^ MEFs and 4T1 induced WTp53 cells. (Fig. S1A-S1C)

The basal activity of the cGAS/STING pathway is due to the presence of endogenous cytoplasmic DNA, and transfection of sheared Herring Testes DNA (HT-DNA) can augment pathway activation. To determine if WTp53 also contributed to the response to HT-DNA, we transfected MEFs (p53^+/+^ vs. p53^−/−^), H1299 (with inducible WTp53), 4T1 (with inducible WTp53) and A549 (shControl vs. shp53) with HT-DNA and assessed the impact on the cGAS/STING pathway. In MEFs, we observed that HT-DNA induced more pronounced phosphorylation of TBK1 substrates in p53^+/+^ MEFs as compared to p53^−/−^ MEFs. (Fig. 1F) Similarly, in WTp53 inducible H1299 cells, we observed a modest induction of TBK1 substrate phosphorylation in HT-DNA transfected uninduced cells, and this was markedly increased in cells induced to express WTp53. (Fig. 1G) Increased phosphorylation of TBK1 substrates was also observed in 4T1 cells in which we induced WTp53 and treated with HT-DNA. (Fig. S1D) Conversely, shRNA knockdown of p53 in A549 cells strongly reduced HT-DNA activation of the cGAS/STING pathway. (Fig. 1E) Assessment of 5 different siRNAs targeting p53 confirmed that p53 knockdown in A549 cells resulted in reduced TBK1 substrate phosphorylation, thus ruling out an off-target effects of the shRNA. (Fig. S1F) Our results indicate that WTp53 can stimulate basal and augment agonist activation of the cGAS/STING pathway.

To determine if increased TBK1 activity in response to WTp53 expression resulted in activation of IRF3’s transcriptional activity, we performed RT-PCR analysis of its canonical transcriptional target, interferon beta 1 (IFNB1). WTp53 induction in H1299 and 4T1 cells increased basal levels of IFNB1 mRNA and also caused a higher induction in response to HT-DNA treatment. (Fig. 1H) Conversely, shRNA knockdown of p53 in A549 cells reduced both the basal expression and HT-DNA induction of IFNB1. (Fig. 1H) In agreement with the transcriptional induction of IFNB1, we observed higher amounts of secreted IFNB1 in cells induced to express p53 (H1299 and 4T1), and reduced IFNB1 secretion in response to p53 knockdown (A549). (Fig. 1I) These results indicate that WTp53 mediated stimulation of the cGAS/STING pathway results in IRF3’s functional activation.

Inactive IRF3 resides in the cytoplasm and upon its phosphorylation by TBK1, translocate to the nucleus.^33,34^ To corroborate our findings that WTp53 activates IRF3, we assessed its subcellular localization. In unstimulated cells, we observed that GFP-tagged IRF3 was primarily located in the cytoplasm. (Fig. S1G) Induction of WTp53 alone was sufficient to promote approximately 20% of cells to exhibit nuclear accumulation of GFP-IRF3. HT-DNA treatment induced GFP-IRF3 nuclear translocation in 50% of the uninduced cells and this was further increased to 80% upon WTp53 induction (Fig. S1G). As a parallel approach to examine IRF3’s localization, we performed subcellular fractionation and detected IRF3 by western blot. In both H1299 and 4T1, WTp53 induction resulted in increased phospho- and total IRF3 accumulation in the nuclear fraction (Fig. S1H). Collectively, our data strongly suggests that WTp53 induces basal and agonist mediated innate immune response.

### WTp53 promotes cGAS/STING/IRF3 mediated apoptosis

Previously, it was reported that potent activation of STING results in apoptosis.^32^ Since we observed that WTp53 could not only stimulate basal but also augment HT-DNA mediated cGAS/STING signaling, we performed FACS analysis to determine if it also dictated the cellular response to activation of the pathway. In uninduced H1299 cells, we observed that HT-DNA transfection induced apoptosis in a dose dependent manner, which further increased in cells induced to express WTp53. (Fig. 2A) Of note, we utilized a WTp53 cDNA that carries a proline at codon 72, a polymorphism that has been shown to minimally induce apoptosis.^35^ In agreement, we observed that WTp53 expression alone did not induce apoptosis. (Fig. 2A) Furthermore, we also treated A549 shControl or shp53 cells with HT-DNA and found that p53 knockdown blunted the apoptotic response to HT-DNA treatment. (Fig. 2B) Thus, our data suggests that WTp53’s potent stimulation of cGAS/STING signaling surpasses the threshold required for apoptosis to occur.

**Figure 2:**
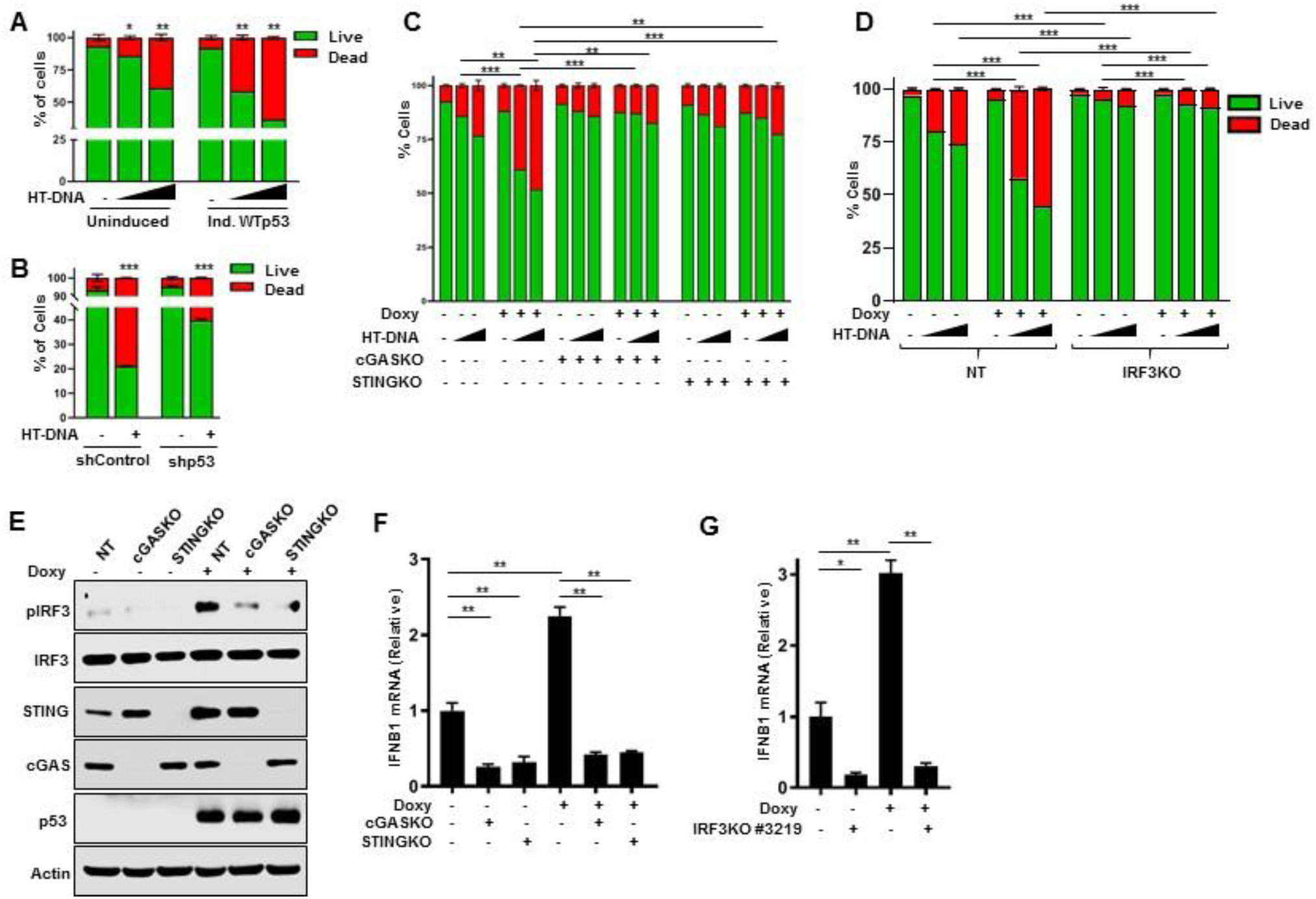
WTp53 promotes cGAS/STING/IRF3 mediated apoptosis. Graphical representation of apoptosis quantification by flow cytometry of (A) H1299 induced WTp53 and (B) shControl or shp53 A549 cells that were treated with HT-DNA for 24 h. H1299 doxycycline inducible WTp53 cells were stably knockout for cGAS, STING and IRF3. Representative graphs indicate quantification analysis of apoptotic death analyzed by flow cytometry of (C) non-target (NT), cGASKO and STINGKO (D) non-target (NT) and IRF3KO H1299 cells induced WTp53 treated with 2 μg/ml or 4 μg/ml of HT-DNA for 24 h. Cells were harvested, stained with Annexin V-FITC and PI and subjected to flow cytometry analysis. (E-F) Non-target (NT), cGASKO and STINGKO H1299 cells induced WTp53 and subjected to (E) Western blot (F) RT-PCR analysis. (G) Non-target (NT) and IRF3KO H1299 cells induced WTp53 cells were subjected to RT-PCR analysis. Quantification graphs: In all panels, error bars represent mean with standard deviation. p values are based on Student’s t test. ***p < 0.001, **p < 0.01, ns=non-significant. See also Figure S2

Thus, we sought to determine if the enhanced apoptotic response in WTp53 induced cells treated with HT-DNA was indeed due to activation of the cGAS/STING pathway. We used CRISPR-Cas9 to knockout cGAS, STING and IRF3 and tested how loss of these genes affected WTp53’s sensitization to HT-DNA. As expected, cGAS knockout reduced the population of cells undergoing apoptosis in response to HT-DNA. Importantly, cGAS knockout completely abrogated the ability of WTp53 to sensitize to HT-DNA treatment.(Fig. 2C) Additionally, both STING and IRF3 knockout cells also rescued the cells from WTp53 sensitization to activation of the pathway. (Fig. 2C and 2D) The p53 transcriptional target IFI16 has been postulated to be a cytosolic DNA sensor and thus we assessed if it was involved in this context. We found that unlike cGAS, IFI16 is dispensable for WTp53 activation of the cGAS/STING pathway and sensitization to HT-DNA. (Fig. S2A) Our data indicates that WTp53 specifically sensitizes to HT-DNA via stimulation of the cGAS/STING pathway.

Since we observed that WTp53 stimulated IFNB1 mRNA expression, and WTp53 has previously been shown to indirectly regulate IFNB1 expression through IRF7 and IRF9, we checked if this was dependent on the cGAS/STING pathway.^12^ Loss of either cGAS or STING resulted in reduced IFNB1 mRNA levels and abrogated WTp53 induction of IFNB1 mRNA. (Fig. 2E and 2F) The involvement of this pathway was further supported by the observation that IRF3 knockout cells also failed to induce IFNB1 mRNA in response to WTp53. (Fig. 2G and S2B) In contrast, IFI16 is not required for WTp53 to stimulate the cGAS/STING pathway since IFI16 knockout had no effect on the stimulation TBK1 substrate phosphorylation or the induction of IFNB1 mRNA. (Fig. S2C and S2D) Taken together our data suggests that WTp53 mediated induction of Type I interferon response is cGAS, STING and IRF3 dependent.

### WTp53 promotes TREX1 degradation resulting in cytosolic DNA accumulation

Our western blot analysis did not indicate that p53 was stimulating the pathway by inducing expression of the components of the cGAS/STING pathway and thus we assessed if it was regulating the activity of cGAS. In both H1299 and 4T1, WTp53 induction resulted in increased cGAMP, the second messenger produced upon agonist activation of cGAS.^30^ (Fig. 3A and 3B) We speculated that the activation of cGAS enzymatic activity by WTp53 is likely due to an increased presence of an endogenous cGAS agonist and thus we assessed the levels of cytoplasmic DNA using either an anti-dsDNA antibody or PicoGreen staining. In A549 cells, we detected cytoplasmic DNA, which decreased after p53 knockdown. (Fig. 3C) Similarly, detection of cytoplasmic DNA by PicoGreen staining exhibited a similar trend of reduced cytoplasmic DNA after p53 knockdown. (Fig. 3D) Conversely, WTp53 expression in H1299 cells resulted in increased detection of cytoplasmic DNA by anti-dsDNA and Picogreen staining. (Fig. 3E and 3F) We also observed that WTp53 induction in 4T1 cells increased the level of cytoplasmic DNA detected with PicoGreen staining. (Fig. S3A) These results suggest that WTp53 stimulates cGAS activity by increasing the abundance of cytoplasmic dsDNA.

**Figure 3:**
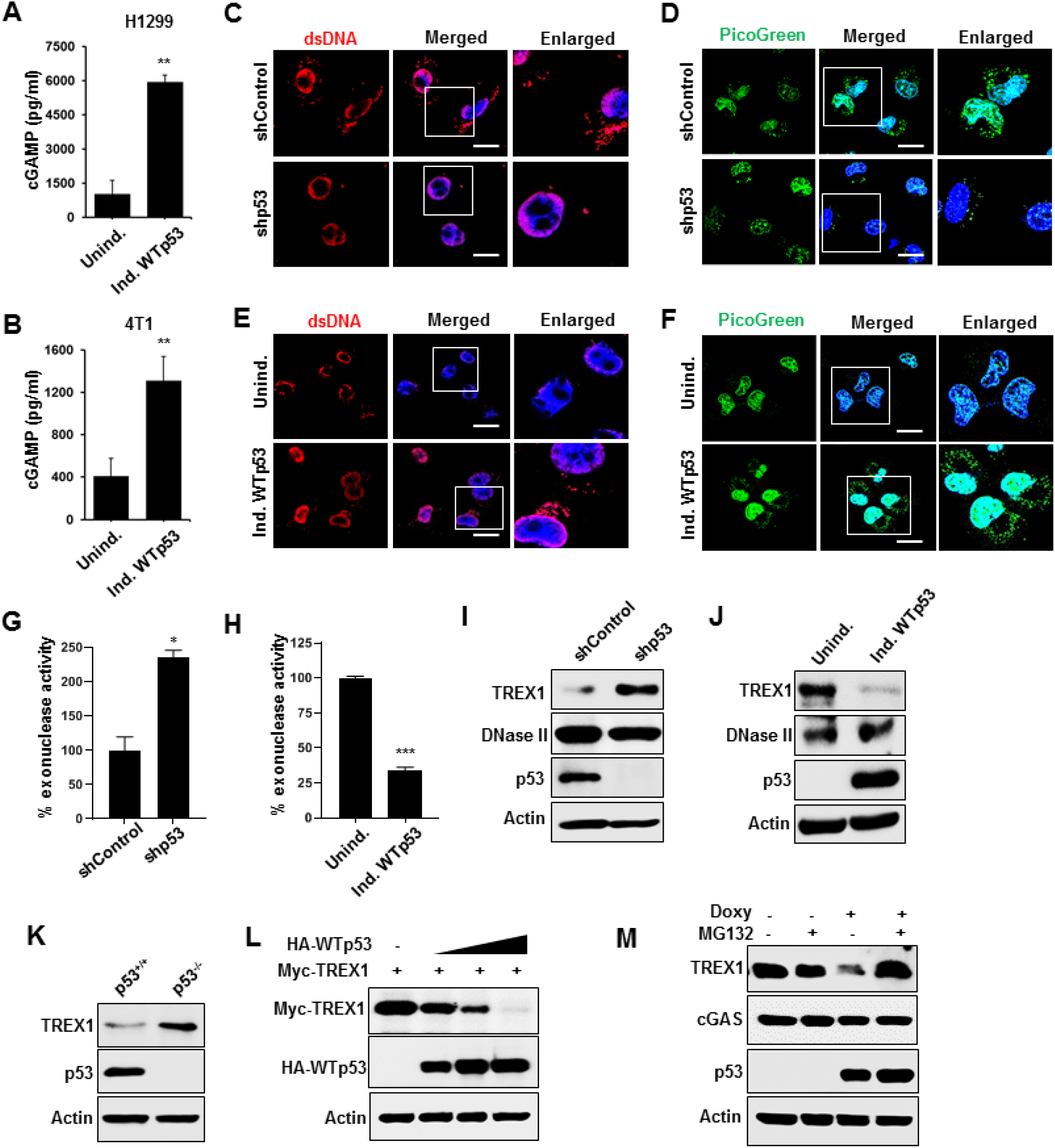
WTp53 promotes TREX1 degradation resulting in cytosolic DNA accumulation. Representative graphs indicate cellular cGAMP levels in (A) H1299 and (B) 4T1 cells with inducible WTp53. (C) Representative confocal microscopy images of cytosolic dsDNA in A549 shControl or shp53 cells with immunocytochemical detection with a dsDNA specific antibody or (D) Confocal live-cell imaging of A549 shControl or shp53 cells stained with 3 μl/ml PicoGreen (green) for 1 hr and counter stained with Hoechst33342 for 15 min. Scale Bar 10 μm. (E) Representative confocal images of H1299 cells expressing inducible WTp53 stained with dsDNA specific antibody. Scale Bar 10 μm. (F) Representative confocal live-cell imaging of H1299 induced WTp53 cells stained with 3 μl/ml PicoGreen (green) for 1 hr and counter stained with Hoechst33342 for 15 min. Scale Bar 10 μm. (G-H) Representative graphs indicates % exonuclease activity in (G) A549 (shControl or shp53) (H) H1299 inducible WTp53 cells. (I) A549 shControl or shp53 cells were subjected to western blot. (J-K) Representative Immunoblots of (J) H1299 cells induce WTp53 and (K) p53+/+ and p53-/- mouse embryonic fibroblast (MEFs). (L) In H1299 cells, Myc-TREX1 was co-transfected with increasing amount of HA-WTp53 and subjected to western blot analysis. (M) H1299 cells were induced with doxycycline to express WTp53 for 24 hrs and treated with MG132 (20 μM) for 6 hrs after which cells were harvested for western blot analysis. Quantification graphs: In all panels, error bars represent mean with standard deviation. p values are based on Student’s t test. ***p < 0.001, **p < 0.01, ns=non-significant. See also Figure S3.

Since cytoplasmic dsDNA is degraded by exonucleases, we assessed if WTp53 regulated cellular exonuclease activity. WTp53 knockdown in A549 cells resulted in an approximately 2.5 fold increase in exonuclease activity, whereas WTp53 induction in H1299 cells resulted in an almost 4 fold decrease. (Fig. 3G and 3H) Similarly, WTp53 induction in 4T1 cells resulted a reduction in exonuclease activity. (Fig. S3B) The 3′→5′ cytosolic exonuclease, TREX1, is the predominant exonuclease, accounting 60 to 70% of total cellular exonuclease activity and thus we assessed its expression.^18,36–38^ TREX1 protein levels were induced after p53 knockdown in A549 and CT26 cells, and reduced in H1299 and 4T1 cells upon p53 induction. (Fig. 3I, 3J and S3C, S3D) Comparison of p53^+/+^ vs. p53^−/−^ MEFs showed increased TREX1 protein in the latter. (Fig. 3K) Analysis of TREX1 mRNA levels in A549 and H1299 cells indicated that WTp53 did not alter its expression, suggesting that it instead controls TREX1 protein turnover. (Fig. S3E) We performed cyclohexamide chase experiments to determine if this was the case. The half-life of the GFP-TREX1 protein in uninduced H1299 cells was approximately 10.8 hours, whereas in WTp53 induced cells it was reduced to 5.5 hours, thus indicating a faster turnover. (Fig. S3F) We co-transfected Myc-TREX1 with increasing amounts of HA-WTp53 and observed a dose dependent reduction in Myc-TREX1. (Fig. 3L) The reduction in endogenous TREX1 levels was rescued by MG132 treatment, indicating that WTp53 promotes TREX1 degradation through the proteasomal pathway. (Fig. 3M) These results suggests that WTp53 induces the accumulation of cytoplasmic dsDNA by promoting TREX1 degradation.

### TRIM24 is an ubiquitin ligase for TREX1

To date, an E3 ubiquitin ligase that promotes TREX1 degradation has not been identified. We speculated that a known p53 transcriptional target with E3 ubiquitin ligase activity might be involved in TREX1 degradation. There are 6 well studied E3 ubiquitin ligases that are p53 transcriptional targets: MDM2, COP1, PIRH2, TRIM24, FBXW7 and SIAH. We transfected H1299 cells with siRNAs targeting these ubiquitin ligases and found that TRIM24 knockdown potently induced TREX1 protein. (Fig. S4A) We also observed a modest induction in TREX1 after FBXW7 knockdown but it also significantly induced TREX1 mRNA. In contrast, TRIM24 knockdown only induced TREX1 protein and had no effect on its mRNA and thus we further assessed if TRIM24 controlled TREX1 turnover. (Fig. S3B) We confirmed in A549 cells that siRNA mediated TRIM24 knockdown induces TREX1. (Fig. 4A) Co-transfection of Myc-TREX1 with Flag-TRIM24 resulted in reduced Myc-TREX1 protein that could be rescued by MG132, indicating that TRIM24 promotes TREX1 degradation through the proteasome. (Fig. 4B) TRIM24’s ability to promote TREX1 degradation required the ring domain since its deletion prevented TREX1 degradation. (Fig. S4C) Additionally, co-immunoprecipitation studies showed that Flag-TRIM24 co-precipitated with GFP-TREX1, suggesting that TRIM24 directly interacts with TREX1 to promote its degradation. (Fig. 4C) Next we performed ubiquitination assays to determine if TRIM24 can promote TREX1 ubiquitination. Transfection of GFP-TREX1 alone resulted in a modest amount of ubiquitination and this was greatly enhanced by co-transfection of Flag-TRIM24. (Fig. 4D) To determine if WTp53 requires TRIM24 to induce TREX1 degradation, we used siRNA to knockdown TRIM24 in H1299 cells transfected with empty vector or a HA-p53 expression vector. We observed that TREX1 downregulation by p53 could be rescued by TRIM24 knockdown, and thus we concluded that TRIM24 mediates TREX1 degradation in response to WTp53. (Fig. S4D)

**Figure 4:**
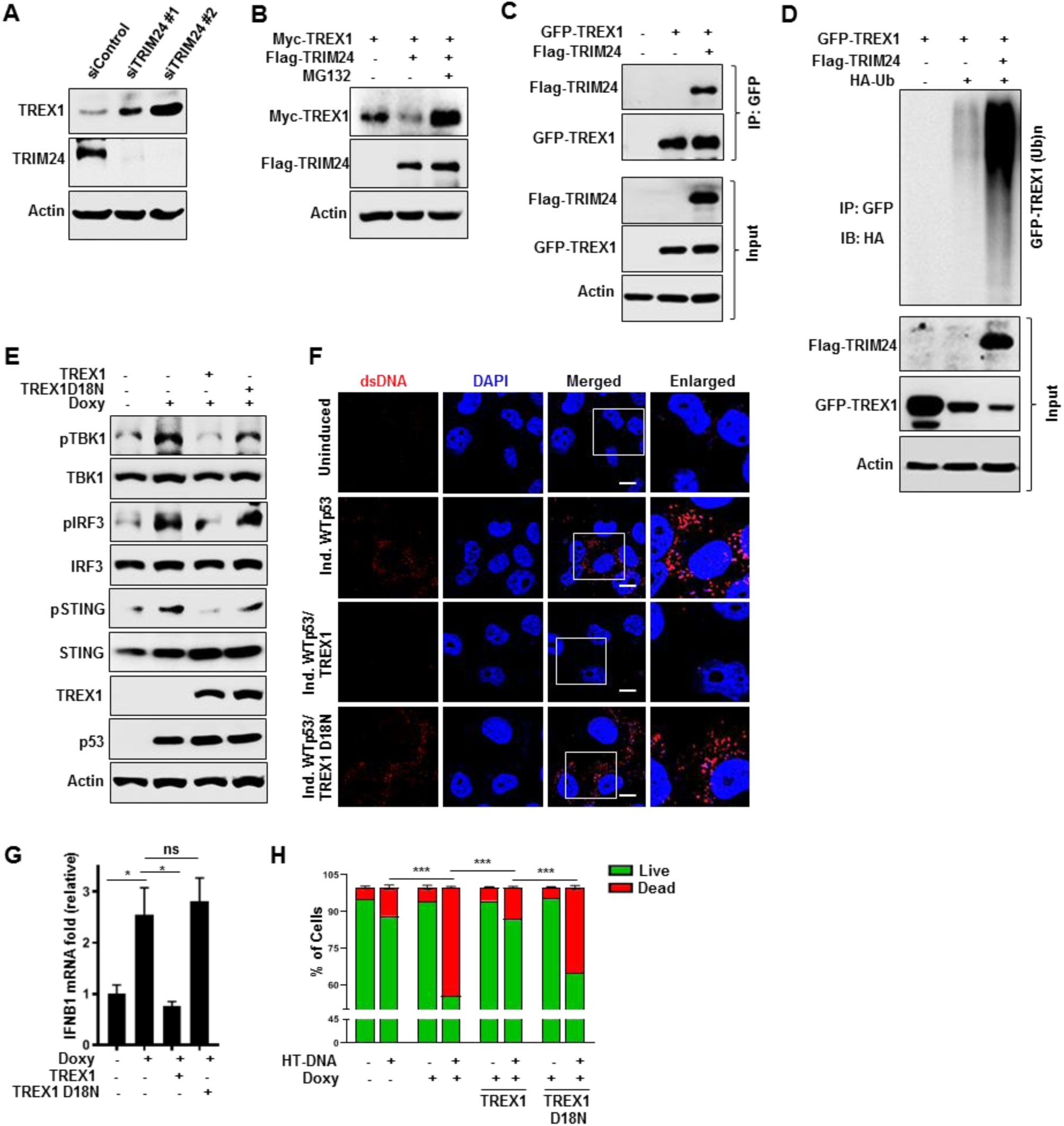
TRIM24 is an ubiquitin ligase for TREX1. (A) A549 cells were transfected with two different siRNA for TRIM24 and subjected to western blot analysis. (B) H1299 cells were co-transfected with Myc-TREX1 and Flag-TRIM24 for 24 h. and prior to harvesting cells were treated with MG-132 for 6 h. (C) p53 null H1299 cells were co-transfected with GFP-TREX1 and Flag-TRIM24. Prior to harvesting, cells were treated with MG-132 for 6 h. Cells were lysed and GFP-TREX1 was immunoprecipitated from the whole cell lysate. Lysates and immunoprecipitates (IP) were analyzed by western blotting. (D) H1299 cells were co-transfected with GFP-TREX1, poly ubiquitin and Flag-TRIM24 for 24 h. Prior to harvesting, cells were treated with MG-132 for 6 h. Cells were harvested under denaturing conditions by using boiling 1% SDS buffer and lysates were then immunoprecipitated and were processed for Western blotting. (E) H1299 induced WTp53 cells were stably overexpress GFP-TREX1 or GFP-TREX1 D18N. Cells were induced to express WTp53 and subjected to western blot. (F) Representative confocal microscopic immunofluorescence (IF) images are showing cytosolic dsDNA in H1299 induced WTp53 overexpress GFP-TREX1 or GFP-TREX1 D18N. Scale Bar 10 μm. (G) H1299 induced WTp53 stably overexpress GFP-TREX1 or GFP-TREX1 D18N cells were lysed and subjected to RT-PCR analysis. (H) Graphical representation of apoptosis quantification by flow cytometry of H1299 induced WTp53 cells were stably overexpress GFP-TREX1 or GFP-TREX1 D18N cells that were treated with 2ug/ml of HT-DNA for 24 hrs. Quantification graphs: In all panels, error bars represent mean with standard deviation. p values are based on Student’s t test. ***p < 0.001, **p < 0.01, ns=non-significant. See also Figure S4.

Our data thus far implicate downregulation of TREX1 protein and exonuclease activity as the mechanism by which WTp53 activates the cGAS/STING pathway and sensitizes cells to agonist induced apoptosis. To test this directly, we co-expressed WTp53 with wildtype TREX1 or its catalytically dead mutant TREX1 D18N, and assessed TBK1 substrate phosphorylation and the accumulation of cytoplasmic DNA. We observed that wildtype TREX1, but not the inactive TREX1 D18N mutant, was able to block the phosphorylation of TBK1 substrates in response to WTp53 expression. (Fig. 4E) Furthermore, wildtype TREX1 overexpression reduced the appearance of cytoplasmic DNA in response to WTp53 and this was dependent on its catalytic activity. (Fig. 4F) Furthermore, wildtype TREX1 (and not the D18N mutant) suppressed the induction of IFNB1 mRNA by WTp53. (Fig. 4G) The observation that WTp53 could augment the activation of cGAS/STING resulting in apoptosis, led us to determine if TREX1 could prevent this from occurring. Expression of WTp53 and treatment with HT-DNA resulted in approximately 55% of the cells undergoing apoptosis. Consistent with our observations that TREX1 could block the activation of the cGAS/STING pathway by WTp53, we observed that TREX1 overexpression suppressed WTp53’s ability to sensitize to HT-DNA. (Fig. 4H) The protection afforded by TREX1 overexpression required its catalytic activity since the D18N mutant was unable to prevent apoptosis. Our data indicate that downregulation of TREX1 by WTp53 augments cGAS/STING pathway activation and sensitizes to innate immune signaling induced apoptosis.

Pharmacological activation of p53 triggers the de-repression of endogenous retroviruses and this has been proposed to trigger interferon expression through a putative mechanism involving PAMP RNA sensing machinery.^11^ We observed that nutlin treatment stimulated phosphorylation of TBK1 and the downregulation of TREX1 in a dose dependent manner. (Fig. S4E) The WTp53 transcriptional target MAVS is a mitochondrial protein that is a downstream effector for various PAMP RNA sensing machinery, and can activate TBK1 and interferon expression.^43–45^ To determine if WTp53 stimulated TBK1 through a PAMP RNA sensing mechanism in a MAVS-dependent manner, we generated MAVS knockout cells with CRISPR/Cas9. Deletion of MAVS had a minimal impact on the activation of TBK1 by WTp53. (Fig. S4F) Our data reinforce the notion that accumulation of cytoplasmic DNA leaked from the mitochondria underlies the activation of the cGAS/STING pathway by WTp53.

### WTp53 promotes cGAS/STING dependent antitumor immune response

Our data led us to posit that WTp53 works through cGAS to regulate tumor growth by recruitment of immune cells. To test this in another model, we generated 4T1 WTp53 inducible cells with a non-targeting or cGAS-targeting CRISPR vector and determined the effect of cGAS loss on WTp53’s control of tumor growth. Consistent with our results showing that cGAS is required for p53 to activate the pathway and induce IFNB1, we observed that WTp53 induction did not induce TBK1 substrate phosphorylation or IFNB1 mRNA in cGAS KO 4T1 cells. (Fig. 5A and 5B) We injected 4T1 inducible WTp53/cGAS knockout cells into the mammary fat pad of female BALB/c mice. We observed that cGASKO tumors grew faster than the control, uninduced tumors. In contrast, doxycycline induction of WTp53 expression resulted in a sustained lack of tumor growth. Surprisingly, WTp53 induced growth suppression was dependent on cGAS since after an approximately 3-day delay in growth, the WTp53-induced/cGASKO tumors resumed growing. (Fig. 5C, S5A and S5B) Immunophenotyping analysis revealed that the blunted growth of the WTp53 induced tumors correlated with a dramatic increase in tumor infiltration by CD45+, CD4+, CD8+ and NK cells. (Fig. 5E-5H) In WTp53-induced/cGASKO tumors, infiltration by these types of immune cells was similar to control tumors, suggesting that cGAS is required for WTp53 to recruit these cells.

**Figure 5:**
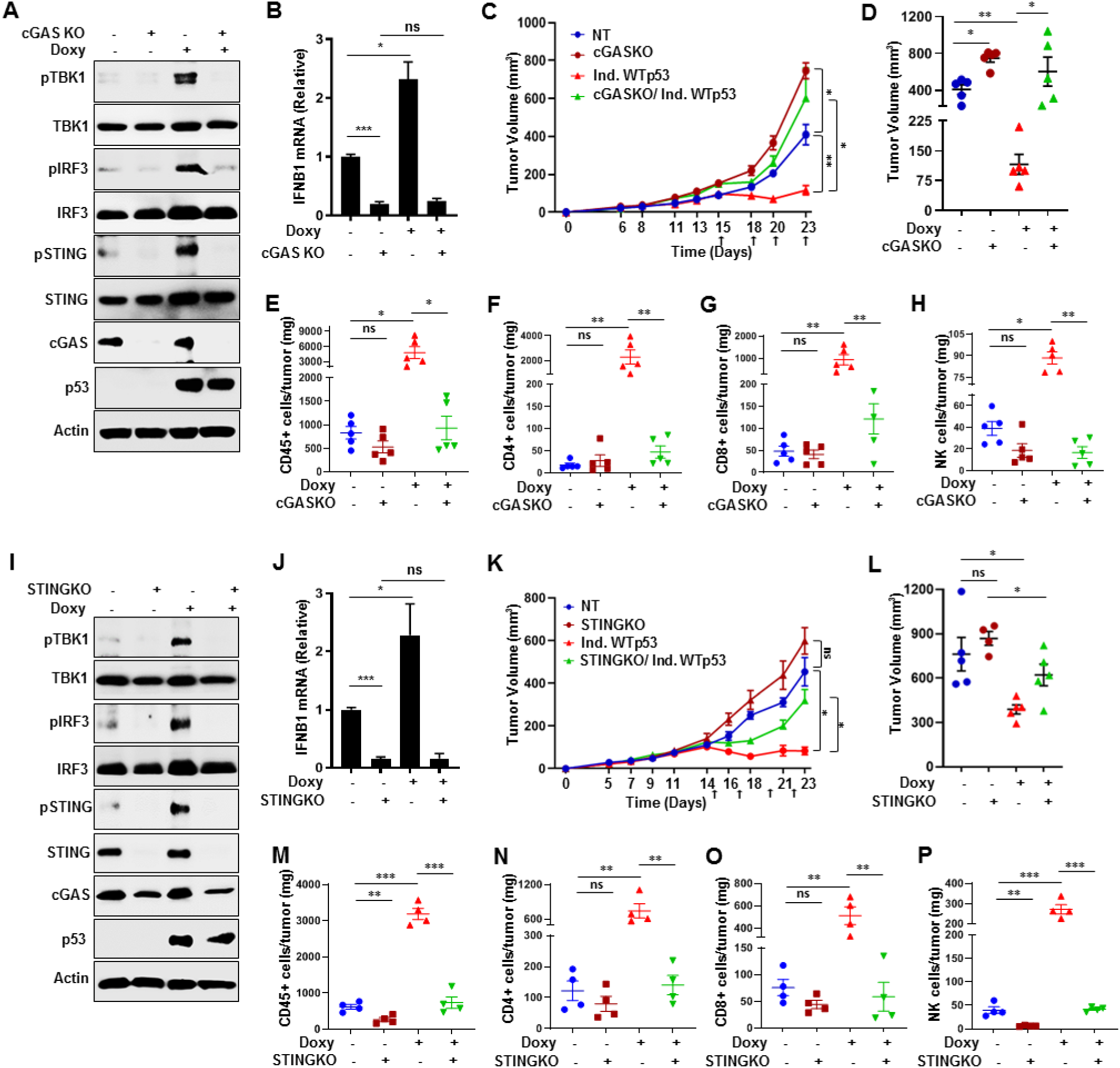
WTp53 promotes cGAS/STING dependent antitumor immune response. (A) Immunoblots of 4T1 doxycycline inducible WTp53 non-target (NT) or cGAS knockout cells. (B) RT-PCR analysis for IFNB1 in 4T1 doxycycline inducible WTp53 non-target (NT) or cGAS knockout cells. (C) 4T1 inducible WTp53 non-target or cGASKO cells were injected into the mammary gland of immunocompetent female BALB/c mice (n=5). Mice were given doxycycline orally to induce WTp53 on the indicated days. Tumor growth was monitored and measured using slide calipers. All mice were sacrificed on day 23 and graphical quantification represents the tumor growth rate. (D) Tumor volume of the indicated 4T1 tumor cohorts on Day 23. (E-H) Representative graphs showed FACS quantification of (E) CD45+ T-lymphocytes (F) CD3+CD4+ T-helper (G) CD3+CD8+ cytotoxic T-lymphocyte and (H) NK cells per milligram (mg) of 4T1 induced WTp53 non-target or cGASKO tumor tissues (n=5). 4T1 doxycycline inducible WTp53 non-target (NT) or STING knockout cells were subjected to (I) western blot or (J) RT-PCR for IFNB1. (K) 5 × 10^4^ 4T1 inducible WTp53 non-target (NT) or STINGKO cells were injected into the mammary gland of immunocompetent female BALB/c mice (n=4). Mice were given doxycycline orally to induce WTp53 on the indicated days (upward arrows). Tumor growth was monitored and measured using slide calipers. All mice were sacrificed on day 23 and graphical quantification represents the tumor growth rate in BALB/c mice. (L) Tumor volume of the indicated 4T1 tumor cohorts on Day 23. (M-P) Representative graphs showed FACS quantification of (M) CD45+ T-lymphocytes (N) CD3+CD4+ T-helper (O) CD3+CD8+ cytotoxic T-lymphocyte and (P) NK cells per milligram (mg) of 4T1 induced WTp53 non-target or STINGKO tumor tissues (n=4). Quantification graphs: In all panels, error bars represent mean with standard error mean. In scatter dot plots, each dot represent one mouse, p values are based on Student’s t test. ***p < 0.001, **p < 0.01, *p < 0.05, ns=non-significant. See also Figure S5.

To further assess the contribution of the immune system to WTp53 growth suppression *in vivo*, we repeated the experimental approach using NOD/SCID mice. In these immunocompromised mice, the growth advantage observed with the cGASKO cells was largely eliminated, indicating that the immune response was required for the observed differences in tumor growth in the syngeneic model. (Fig. S5C-S5E) WTp53 induction slowed tumor growth as compared to the other groups, however, unlike in the BALB/c mice, the tumors continued to grow and did not plateau. The lack of sustained growth arrest suggests that the immune system has a key role in p53-mediated growth suppression. Importantly, cGASKO again rescued the reduced tumor growth of the WTp53 induced tumors. These data suggest that cGAS is a downstream effector of WTp53’s growth inhibitory function through both cell-autonomous and non-cell autonomous mechanisms.

cGAS has been show to regulate cellular phenotypes through STING-dependent and independent mechanisms.^54^ The pronounced requirement for cGAS in WTp53 tumor suppression led us to investigate if STING was similarly involved. We used CRISPR/Cas9 to generate WTp53 inducible 4T1 cells that lack STING (STINGKO). Similar to the cGASKO data above, STINGKO cells failed to induce TBK1 substrate phosphorylation or IFNB1 mRNA in response to WTp53 expression. (Fig. 5I, 5J) As expected, we observed that WTp53 reduced proliferation *in vitro* and that STINGKO cells had a slightly higher proliferation rate but it was not statistically significant different from control cells. (Fig. S5F) When injected into syngeneic mice, the STINGKO tumors grew slightly faster than the control tumors but the difference was not statistically significant. (Fig. 5K, 5L) As before, we observed that WTp53 expression induced a stable suppression of tumor growth. Strikingly, STING knockout rescued WTp53 induced tumor growth suppression. (Fig. 5K, S5G and S5H) In keeping with a role for the cGAS/STING pathway in the recruitment of immune cells by WTp53, we observed that STING KO prevented the spike in TILs in the WTp53-induced/STINGKO tumors. (Fig. 5M-5P)

To further assess the immune system involvement in WTp53 mediated tumor growth suppression *in vivo*, we injected 4T1 STINGKO inducible WTp53 cells into immunodeficient NOD/SCID mice. Whereas WTp53 induced tumors grew slower, STING KO tumors grew faster but not in a statistically significant manner compared to the non-targeted control tumors. (Fig. S5I-S5K) STINGKO partially rescued WTp53 mediated tumor growth suppression suggesting that like cGAS, STING also is a downstream effector of WTp53’s growth inhibitory function. Taken together, our study reveals that WTp53 engagement of the cGAS/STING pathway is critical for its tumor suppressor activities through tumor cell autonomous and non-cell-autonomous mechanisms.

### TREX1 deficiency induces a WTp53 dependent antitumor immune response

Genetic perturbation of TREX1 function results in a severe autoimmune response due to the cellular inability to clear cytosolic DNA.^19,46–49^ This autoimmune response due to TREX1 deficiency can be rescued by loss of either cGAS, STING or IRF3, indicating that these genes are epistatic.^19,48,50–53^ Since WTp53 positively regulates the cGAS/STING pathway, we considered the possibility that WTp53 knockdown may also negate the immunostimulatory effect of TREX1 loss. We used CT26 cells to test this possibility *in vivo* in a syngeneic BALB/c mouse tumor model. We generated CT26 cells with shRNA mediated knockdown of either p53, TREX1 alone or both. (Fig. S6A) Western blot analysis of these different shRNA knockdown cells showed a decrease in cGAS/STING pathway upon p53 knockdown. As expected, cells with TREX1 knockdown had elevated activity of the cGAS/STING pathway as indicated by the increased phosphorylation of TBK1 substrates and higher IFNB1 mRNA expression. The combined knockdown of p53 and TREX1 reverted the cGAS/STING pathway activity and IFNB1 mRNA back to baseline control levels. (Fig. S6B) Next, we injected CT26 shControl, shp53, shTREX1 and shp53/shTREX1 cells subcutaneously in BALB/c mice and monitored tumor growth. The hyperactivation of the cGAS/STING pathway resulted in a severe growth delay in the TREX1 knockdown tumors. (Fig. 6A) In contrast, p53 knockdown cells produced tumors that grew faster than the control. Strikingly, the combined knockdown of TREX1 and p53 resulted in tumors that grew at the same rate as the control tumors. (Fig. 6A and S6C, S6D) The observed differences in tumor growth *in vivo* did not reflect differences in cell proliferation as we did not observe any significant differences *in vitro*. (Fig. S6E) Additionally, when these same cells were injected into immunocompromised NOD/SCID mice, there was no difference in tumor growth, suggesting that an intact immune system was required to differentially control tumor growth. (Fig. 6B and S6F, S6G) To gain insight into this observation we performed immune-phenotyping to detect the presence of tumor infiltrating immune cells (Fig. S6H). WTp53 knockdown reduced the intratumoral presence of CD45+, CD4+, CD8+ and NK cells. In contrast, TREX1 knockdown tumors had a completely opposite phenotype and showed massive infiltration by those immune cells types. Importantly, the combined knockdown of TREX1 and p53 resulted in an immune infiltration phenotype similar to control tumors. (Fig. 6C-6F) Thus, p53 loss is similar to cGAS/STING/IRF3 loss and is sufficient to negate the effect of TREX1 deficiency.

**Figure 6:**
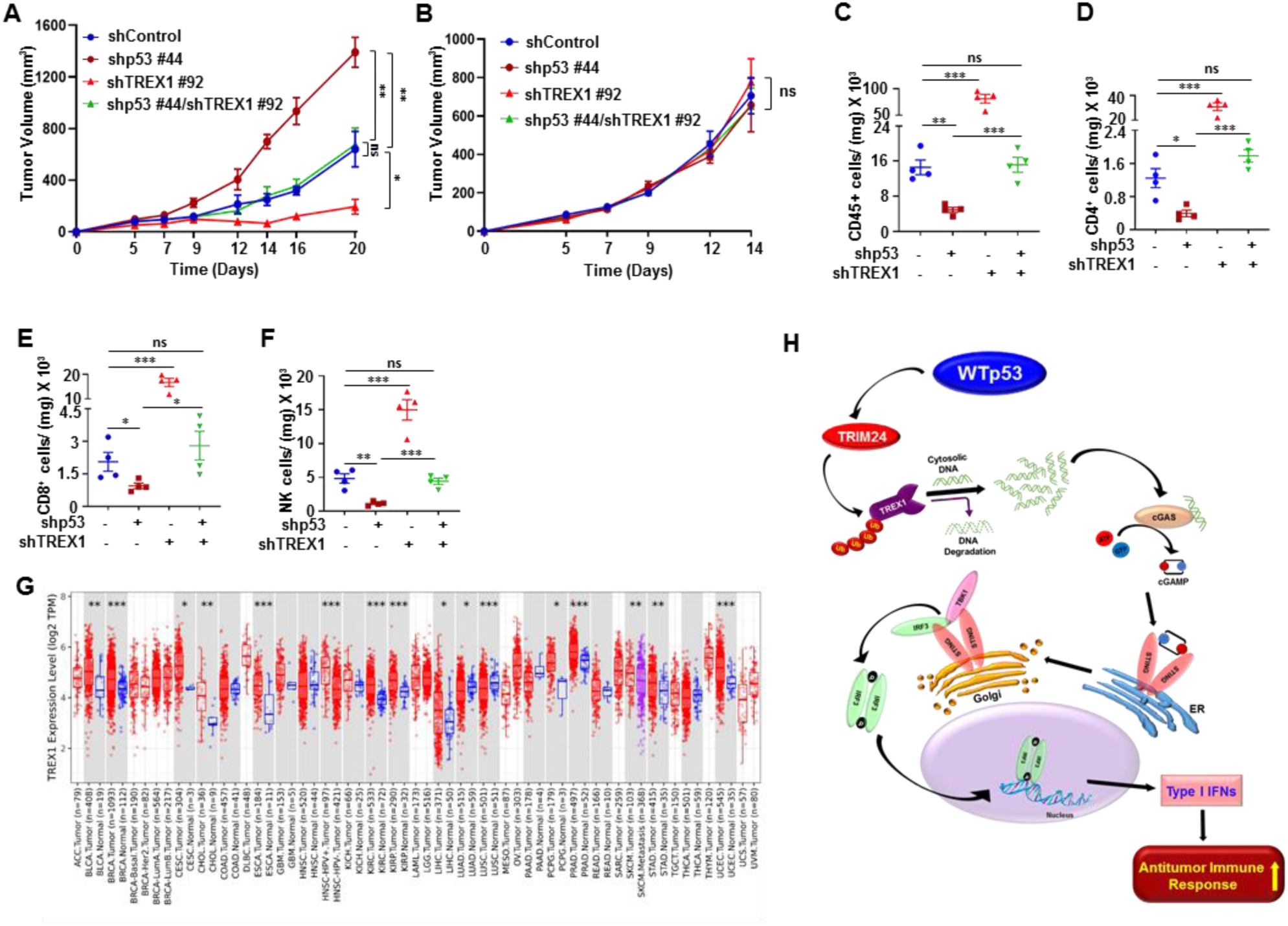
TREX1 deficiency induces a WTp53 dependent antitumor immune response. (A) CT26 shControl, shp53, shTREX1 and shp53/shTREX1 cells were injected with matrigel subcutaneously into immunocompetent female BALB/c mice (n=4). Tumor growth was monitored and measured using slide calipers. All mice were sacrificed on day 21 and graphical quantification represents the tumor growth rate in mice. (B) CT26 shp53, shTREX1 or shp53/shTREX1 cells were subcutaneously injected into the of immunodeficient NOD/SCID mice (n=5). All mice were sacked on day 14 and graphical quantification represents the tumor growth rate. (C-F) Representative graphs showed FACS quantification of (C) CD45+ T-lymphocytes (D) CD3+CD4+ T-helper (E) CD3+CD8+ cytotoxic T-lymphocyte and (F) NK cells per milligram (mg) of indicated tumor tissues (n=4). (G) TREX1 expression in different tumor tissues in the Tumor Immune Estimation Resource (TIMER) database (http://timer.cistrome.org/). (*p < 0.05, **p < 0.01, ***p < 0.001). (H) Schematic representation of WTp53 promotes TRIM24 mediated TREX1 degradation that cause cytosolic DNA accumulation and activation of innate immune pathway to induce antitumor immune response. Quantification graphs: In all panels, error bars represent mean with standard deviation. p values are based on Student’s t test. ***p < 0.001, **p < 0.01, ns=non-significant. See also Figure S6.

Since we observed that TREX1 could prevent the activation of the cGAS/STING pathway by p53, we surmised that TREX1 might function as an oncogene in human cancers and thus we performed a pan-cancer analysis of its expression using TCGA and GEPIA data sets. In comparison to normal tissue, TREX1 was overexpressed in a variety of cancers. (Figure 6G and S6I). We also analyzed the Human Protein Atlas (HPA) database and found TREX1 was highly overexpressed in the tumor tissue when compared with the normal tissue. (Fig. S6J) Furthermore, when we compared survival between low and high TREX1 expressing breast cancer patients, we observed that high TREX1 expression portended poor long-term survival. (Figure S6K) These data suggest that elevated TREX1 expression contributes to tumor aggressiveness.

## Discussion

The tumor suppressor p53 has been shown to regulate a variety of cellular processes that contribute to its tumor suppressor activity. Although studies have shown that p53 induced crosstalk between tumor cells and the immune system can lead to cancer immunoediting, the underlying mechanism(s) are not well understood. Recently, it was shown that pharmacological activation of p53 with nutlin induced expression of endogenous retroviruses (ERVs) and the accumulation of double stranded RNAs (dsRNA) in the cytoplasm.^11^ The dsRNA was associated with the stimulation of Type I interferons, which was purportedly a result of activation of innate immune response RNA sensors.^11^ However, our data show that MAVS knockout did not prevent activation of TBK1 by WTp53 suggesting that MAVS-dependent RNA sensing machinery is not involved. Importantly, nutlin treatment promoted TREX1 degradation and TBK1 activation, indicating that pharmacological activation of p53 recapitulates the signaling to the cGAS/STING pathway that we observed. Moreover, TREX1 overexpression is sufficient to override the stimulation of the cGAS/STING pathway by WTp53, which lends credence to our interpretation that the key activating event is the accumulation of cytoplasmic DNA resulting from TREX1 degradation. This is further supported by the observation that WTp53 can lower the threshold for HT-DNA activation of the pro-apoptotic activity of the cGAS/STING pathway, and this can be reversed by TREX1 overexpression. Our *in vivo* studies reinforce the notion that WTp53’s regulation of TREX1, cGAS and STING mechanistically drives its ability to modulate immune cell activity. Strikingly, our data also revealed that cGAS and STING play an integral role in WTp53 tumor growth suppression *in vivo*, independently of their effects on the immune system.

We observed that p53 lowered the threshold for apoptosis induction by HT-DNA, a cGAS agonist. Therefore, it is possible that combining pharmacological activators of p53 with cGAS or STING agonists could enhance the immunogenicity of tumor cells and their subsequent elimination. Furthermore, we found that TREX1 overexpression suppressed the activation of the cGAS/STING pathway. Based on the observation that TREX1 is overexpressed in a number of human cancers, our data suggest that the development of TREX1 inhibitors has the potential to restore antitumor immune responses in different types of cancers. Moreover, targeting TREX1 in tumors that have lost wildtype p53 may render them immunologically “hot” allowing for immune cell dependent clearance. Taken together, our study establishes danger-associated molecular pattern (DAMP) sensing as a novel component of WTp53’s tumor suppressor activity.

## Acknowledgments

The authors wish to acknowledge the Stony Brook Cancer Center for expert assistance with the Nanostring Data analysis. We would like to acknowledge the technical support provided by the Research Flow cytometry core facility, Department of Pathology, Stony Brook Renaissance School of Medicine.

## Funding

This work was supported by NCI CA166974-07A1, the New York State Empire Investment Program, the Stony Brook Renaissance School of Medicine and Cancer Center, the TRO Carol M. Baldwin Award, the Stony Brook Cancer Center Bahl IDEA Award, and the Lynn November Pilot Funds for Therapeutic Development in Aggressive Breast Cancer and the John C. Dunphy Private Foundation Cancer Innovation Fund from the Stony Brook Cancer Center. D.C.M. is supported by NIH/NCI K22CA226033, a grant from the American Pulse Association, and startup funds from the Stony Brook Cancer Center (Stony Brook, NY) and Bahl Center for Metabolomics and Imaging (Stony Brook, NY). We thank Dr. Nancy C. Reich (Stony Brook University, NY, USA) for providing us with the pcDNA3 GFP-IRF3 clone.

## Author contributions

L.A.M. conceived and supervised the project. L.A.M. and M.G. designed the research. M.G. performed most of the experiments, L.A.M. assisted in experiments. S.S. helped in all the animal experiments, tumor immune profiling and statistical analysis. L.A.M., M.G., and S.S. analyzed the data. L.A.M. and M.G. wrote the manuscript. D.C.M. helped in animal experiments and edited the manuscript. J.L. analyzed the Nanostring data.

## Competing interests

Authors declare that they have no competing interests.

## Notes

### Competing Interest Statement

The authors have declared no competing interest.

