## Supplementary File for "p53 engages the cGAS/STING cytosolic DNA sensing pathway for tumor suppression"

#### Method

##### Mice

4-6 weeks old male or female BALB/c, NOD/SCID mice were purchased from Envigo. All the experiments with mice were conducted in Stony Brook University animal care facility and in accordance with the Institutional Animal Care and Use Committee.

##### Cell Lines

All the cell lines were purchased from ATCC and cultured according to the manufacturer's instructions. H1299, A549 (Human), 4T1, CT26 (Mouse) cells were cultured in complete Roswell Park Memorial Institute (RPMI) medium supplemented with 10% Fetal Bovine Serum (FBS). HEK293T was cultured in complete Dulbecco's Modified Eagle Medium (DMEM) medium. MEFs ($p53^{-/-}$, $p53^{+/+}$) were isolated in Dr. Tomoo Iwakuma Lab, University of Kansas Medical Center, according to the protocol described earlier and were cultured in complete DMEM medium.[55]

##### Cloning and Plasmids

Human shp53 pLKO.1 was a kind gift from Robert Weinberg (Addgene #19119). Non-targeting control and mouse p53 shRNA and TREX1 shRNA were cloned into pLKO.1 (Addgene #24150 and #8453) vector. Doxycycline inducible WTp53 was cloned in pCW57-MCS1-2A-MCS2 (Addgene # 71782). cGAS, STING, IRF3, IFI16 gRNAs were cloned into lentiCRISPRv2-puro (Addgene # 98290) as previously described.[56] GFP-TREX1 (Addgene #27219) and GFP-TREX1 D18N (Addgene #27220) cDNAs were cloned into the lentivirus vector PLVX-puro.

pCMVHA-WTp53 was generated by PCR amplification of p53 from IMR90 lung fibroblast cDNA and cloned into pCMV-HA (Clontech). pcDNA3-GFP-IRF3 was a kind gift from Nancy Reich (Stony Brook University). All the constructs used in the study were confirmed by DNA sequencing. The sequences for sgRNAs, shRNAs, siRNAs and primers for PCR mutagenesis used in this study are listed in Supplementary Table S1.

**Lentiviral particles production in 293FT cells.**

For shRNA knockdown lentiviral particle was generated by transfecting 293T cells with 1.5 µg of ps-PAX2 (Addgene #12260), 0.5 µg of pCMV-VSV-G (Addgene # 8454) and 2 µg of plasmid of gene of interest using Lipofectamine 2000 following the manufacturer's protocol. Viral supernatant was collected post 48 hrs and 72 hrs of transfection. Target cells were infected with the viral particle using polybrene (5 µg/mL) overnight and selection was performed with puromycin or hygromycin.

**Generation of knock out cells using CRISPR-Cas9**

To generate lentiviruses for transduction, HEK293T cells were transfected with plasmid(s) encoding guide RNAs targeting selected genes and packaging vectors (pCMV-VSV-G and psPAX2) using standard Lipofectamine 2000 transfection method. Culture supernatants were collected at 48 and 72 h post-transfection and used for infection of the targeted cells with polybrene (5 µg/ml). Cells were selected with puromycin for 10 days.

**Cytoplasmic and nuclear fractionation**

Doxycycline inducible WTp53 H1299 and 4T1 cells were culture in 60mm dishes and doxycycline was added for 24 hrs to induce p53. Cells were washed with PBS, harvested with trypsin-EDTA, and washed twice with PBS to remove traces of trypsin and growth medium. Cytoplasmic and nuclear fractionation was performed using the subcellular protein fractionation kit (Thermo Scientific, 78840) according to the manufacturer's instructions.

**Cell proliferation assay.**

CT26 shControl or shp53 cells (3,000) and 4T1 induced WTp53 STINGKO cells (1000) were seeded on a 96-well plate and cell proliferation was detected for the next five days. Viable cells were measured by CellTiter-Blue® Cell Viability Assay kit (Promega) according to the manufacturer's protocol. Briefly, 20 µl of cell titer blue reagent was directly added to the culture medium and incubated at 37°C for 4 h and plates were shaken for 10 sec and the fluorescence reading were obtained by reading the plate at 570/590 nm by Molecular Device Spectra Max M5 instrument.

**Real-time quantitative PCR (RT-PCR).**

Cells were cultured according to the experiment and total RNA were collected in RLT buffer (Qiagen) and isolated using the Qiagen mini RNA isolation kit. RNA quantity and quality were confirmed with a NanoDrop ND-1000 spectrophotometer, cDNA was synthesized using 500 ng of total RNA using oligo (dT) primers and Reverse Transcriptase (Quanta). Real-time qRT-PCR was performed in Bio-Rad CFX96 Touch real-time PCR detection system using Universal SYBr Green Supermix (Bio-Rad). Gene-specific primers sequences are listed in Supplementary Table S1.

**IFN Beta measurement by ELISA.**

H1299 and 4T1 doxycycline inducible WTp53 cells ($10^6$) and A549 shp53 cells ($10^6$) were seeded on a 60 mm dish and after 24 hrs cells were transfected with HT-DNA (4 µg) for 18 hrs and cellular conditioned media were collected analyzed using VeriKine Human or mouse IFN Beta ELISA Kit. Quantification of IFNB1 concentration was performed in triplicates according to the manufacturer protocol and the reading was taken at 450 nm by Molecular Device Spectra Max M5 instrument and calculated using an IFNB1 standard curve.

**Nanostring gene Analysis**

H1299 inducible WTp53 cells were treated with doxycycline to induce WTp53 for 24 hrs and A549 shControl and shp53 cells were harvested and lysed in RLT (Qiagen) buffer. RNA was extracted using the manufacturer's protocol. Samples were prepared according to the manufacturer's protocols for the Nanostring nCounter Autoimmune Profiling Panel (NanoString, Seattle, WA, USA). A list of genes and target probe sequences can be found at www.nanostring.com. Cartridges were run on the nCounter Sprint Profiler. Transcripts were analyzed using the nSolver software and R studio v3.6.

**cGAMP ELISA.**

Cytosolic cGAMP was measured using Cayman Chemical 2′ 3′-cGAMP ELISA Kit according to manufacturer's protocol. For cells quantification, 10 million cells were harvested, washed with PBS, Cells were lysed using lysis buffer from Thermo (as recommended by the Manufacturer). Each cell lysate was examined according to the manufacturer's instructions.

**siRNA mediated transient knockdown**

Cells ($2X10^5$) were plated onto a 6-well plate and after 24 hrs siRNA of selected genes were transfected using Lipofectamine RNAiMAX (Invitrogen) using the manufacturer's protocol. Cells were harvested after 48 hrs and processed for western blot or RT-PCR analysis.

**Immunofluorescence.**

H1299 cells with inducible WTp53 cells stably transfected with GFP-IRF3 were grown onto 1% gelatin pre-coated glass coverslips and after all the treatment cells were washed twice with DPBS and counter stained with Hoechst 33342 and mounted with the ProLong Gold Antifade Reagent. Images were captured with a Nikon T*i* epifluorescence microscope and processed using Nikon AR software. For immunofluorescence, cells were grown on glass cover slides and after all the treatment cells were washed twice with DPBS and fixed with 4% paraformaldehyde for 15 mins at room temperature. Cells were then permeabilized with 0.1% triton X-100 and block in 2% BSA for 45 mins. Cells were then incubated with corresponding primary antibodies overnight at 4°C. Cells were washed twice and incubated with secondary antibody for 2 hrs at room temperature. Secondary antibody washed counterstained by DAPI and mount on slides with Fluoromount G.

Cells were stained with 3 µl/ml PicoGreen for 1 hr at 37°C and counter stained by Hoechst 33342. Staining was examined under a Leica confocal scanning microscope equipped with a 100× oil-immersion objective.

**Mitochondria depleted Rho0 cell preparation**

H1299 inducible WTp53 cells were cultured in tetracycline free RPMI supplemented with 500 ng/ml ethidium bromide (Sigma-Aldrich), 50 µg/ml uridine and 1 mM sodium pyruvate for 12 Days. Mitochondrial depletion was checked under CLSM using Mitotracker Red dye.

**Flow cytometry.**

Cells were seeded in a 6-well plate and after all the treatment cells were washed twice with PBS, trypsinized and re-suspended in 100 µl of binding buffer and further incubated with Annexin-V FITC or Annexin-V Pacific blue and Propidium iodide for 15 min in the dark at room temperature. Prior to flow cytometric analysis, 400 µl of binding buffer was added and cells were immediately subjected for the FACS analysis for the number of apoptotic cells. Data was generated using Cole Parmer Cytoflex.

**Preparation of single cells from tumors for Tumor Immune Profiling**

For analyzing tumor associated immune cell populations, tumors were excised, finely minced and incubated in RPMI 1640 containing 0.5 mg/ml collagenase D (Worthington), 0.01 mg/ml DNase I (Roche), and 0.5 mg/ml Dispase (Worthington) for 30 min at 37°C on a shaking platform. Post digestion, the cells were filtered using 70 µm cell strainer. Samples were pelleted at 1000 rcf for 5 mins at 4C and washed with PBS. The red blood cells (RBCs) of the samples were lysed using ACK lysis buffer (Invitrogen). Samples were washed with FACS buffer (PBS+2% FBS) and subjected to processing for flow cytometric analysis. Samples were incubated with Fc blocking antibody (BioLegend) for 15 mins at room temperature and subjected to live dead staining as well as cell surface marker staining using fluorochrome-conjugated monoclonal antibodies. Antibodies are listed below and were purchased from BioLegend, eBioscience and R&D: CD45 (clone 30-F11), CD3ε (145-2C11), CD4 (RM4.5), CD8 (53-6.7), B220 (RA3-6B2), MHC class II (M5/114.15.2) CD11b (M1/70), CD11c (N418), Ly6G (1A8), F4/80 (BM8), CD206 (MMR). Stained cells were washed with cold PBS +1% FBS and kept on ice until analysis. Gating was based on live cells and live cell populations were discriminated initially via CD45/SSC scatterplots, and the different cell populations were defined based on our gating strategy. Fluorescence minus One (FMO) controls were performed as well. Stained cells were acquired using BD LSRII flow cytometry (BD) and analyzed using the Kalluza software.

**Immunoblotting, Ubiquitination and Immunoprecipitation.**

To prepare cell lysates for western blotting, the cells were lysed on the dish using RIPA (0.5% SDS, 0.1% Sodium Deoxycholate, 0.5% NP40, 1 mM EDTA, in PBS pH 7.4 and filter-sterilize) buffer supplemented with protease and phosphatase inhibitors, scraped and placed into microcentrifuge tubes, sonicated and centrifuged at 13,000g for 10 mins at 4 °C to remove insoluble material. Protein concentration was determined using the Micro BCA Protein Assay kit (Pearce) and equal amounts of protein were resolved on 8 or 10% Bis-Tris polyacrylamide gels, transferred to a PVDF membrane blocked with 5% milk and incubated with primary antibody over night at 4C. For co-immunoprecipitation of proteins, cells were washed with PBS, harvested and lysed in immunoprecipitation buffer (50 mM Tris-HCl pH 8.0, 150 mM NaCl, 0.05 mM EDTA, 1% NP40 and 10% glycerol). Lysate was clarified by centrifugation at 20,000g (4°C) for 20 min, pre-cleared with protein-G agarose (KPL) for 2 h at 4 °C and then immunoprecipitated overnight with the corresponding antibodies.

***In vivo* animal experiments.**

Mice were anesthetized using Isoflurane and CT26 shControl or shp53 cells (50,000) in 0.1 ml PBS were injected with matrigel into mice. Mice were monitored and tumor volume was measured manually using slide calipers every other day till day 21 when all the mice were sacked and the tumors were harvested.

4T1 inducible WTp53 cGAS or STING KO cells were trypsinized washed twice with PBS and 50000 cells in PBS were injected at the mammary gland after anaesthetizing the mice. When the tumors reaches 100 $mm^3$, mice were given doxycycline (20 mg/kg) orally every other day to induce WTp53. Tumors were monitored and volume was measured manually using slide calipers till day 21 when all the mice were sacked and the tumors were processed for further experiment.

**Statistical analysis.**

All the experiments were repeated at least three times unless otherwise mentioned in the figure legends. The statistical differences in all assays including Fold difference in mRNA, cell proliferation and growth, flow cytometry and tumor growth between different samples and/or treatments were analyzed by two-tailed Student's t-tests using Microsoft Excel 2013 and all the graphs were made on GraphPad Prism 8 (GraphPad Software) and presented as Mean ±SD or Mean ±SEM. The gene expression analysis was carried out by NanoString using their nSolver software. Statistical significance was set at $P < 0.05$, unless otherwise stated in the text. All experiments were carried out with at least three biological replicates otherwise mentioned in the figure legend. The numbers of animals used are described in the corresponding figure legends.

**KEY RESOURCES TABLE**

| REAGENT or RESOURCE | SOURCE | IDENTIFIER |
|---|---|---|
| Antibodies | | |
| STING (D2P2F) | Cell Signaling | Cat# 13647, RRID:AB_2732796 |
| TBK1 / NAK | Cell Signaling | Cat# 3504, RRID:AB_2255663 |
| IRF3 | Cell Signaling | Cat# 4302, RRID:AB_1904036 |
| Phospho-STING (Ser366) | Cell Signaling | Cat# 19781, RRID:AB_2737062 |
| Phospho-TBK1 (Ser172) (D52C2) | Cell Signaling | Cat# 5483, RRID:AB_10693472 |
| Phospho-IRF-3 (Ser396) (D6O1M) | Cell Signaling | Cat# 29047, RRID:AB_2773013 |
| TP53 | Santa Cruz Biotechnology | Cat# sc-126, RRID:AB_628082 |
| p53 (CM5) | Leica | Cat# sc-6243, RRID:AB_653753 |
| GAPDH | GeneTex | Cat# GTX100118, RRID:AB_1080976 |
| Anti-beta-Actin-Peroxidase antibody | Sigma-Aldrich | Cat# A3854, RRID:AB_262011 |
| c-Myc | Santa Cruz Biotechnology | Cat# sc-40, RRID:AB_627268 |
| HA-probe (F-7) | Santa Cruz Biotechnology | Cat# sc-7392, RRID:AB_627809 |
| GFP (B-2) | Santa Cruz Biotechnology | Cat# sc-9996, RRID:AB_627695 |
| Flag-HRP | Sigma-Aldrich | Cat # A8592, RRID:AB_439702 |
| anti-mouse CD335 (NKp46) antibody | BioLegend | Cat# 137611, RRID:AB_10915472 |
| anti-mouse CD45 | Biolegend | Cat# 103131, RRID:AB_893344 |
| CD3e Monoclonal Antibody (145-2C11) | Thermo Fisher Scientific | Cat# 25-0031-81, RRID:AB_469571 |

| | | |
|---|---|---|
| CD4 Monoclonal Antibody (RM4-5) | Thermo Fisher Scientific | Cat# 17-0042-81, RRID:AB_469322 |
| anti-mouse CD8a | BioLegend | Cat# 100707, RRID:AB_312746 |
| TREX1 | Abcam | Cat# ab185228, RRID:AB_2885196 |
| TRIM24 | Proteintech | Cat# 14208-1-AP, RRID:AB_2256646 |
| V5-probe (E10) | Santa Cruz Biotechnology | Cat# sc-81594, RRID:AB_1131162 |
| cGAS | Cell Signaling | Cat# 15102, RRID:AB_2732795 |
| Anti-Lamine B1 | Abcam | Cat# ab16048, RRID:AB_443298 |
| dsDNA | Santa Cruz Biotechnology | Cat# sc-58749, RRID:AB_783088 |
| IFI-16 (1G7) | Santa Cruz Biotechnology | Cat# sc-8023, RRID:AB_627775 |
| Chemicals, Peptides, and Recombinant Proteins | | |
| Lipofectamine 2000 | Invitrogen | Cat# 2082816 |
| Lipofectamine RNA iMAX | Invitrogen | Cat# 13778-150 |
| PicoGreen | Invitrogen | Cat# P7589 |
| Cyclosporine A | ApexBio | Cat# B1922 |
| Ethidium bromide | Sigma-Aldrich | Cat# 15585-04 |
| HT-DNA | Sigma-Aldrich | Cat# D6898 |
| Doxycycline hyclate | Sigma-Aldrich | Cat# D9891 |
| Sodium pyruvate | Gibco | Cat# 11360-070 |
| Hoechst 33342 | ThermoFisher Scientific | Cat# H3570 |
| DAPI | ThermoFisher Scientific | Cat# 62248 |
| FITC Annexin V/ PI | BioLegend | Cat# 640914 |
| Doxycycline hydrochloride | Fisher Scientific | Cat# BP2653-5 |
| Halt™ Protease Inhibitor Cocktail | ThermoFisher Scientific | Cat# 78429 |
| Fluoromount G | SuthernBiotech | Cat# 0100-01 |
| ProLong™ Gold Antifade | ThermoFisher Scientific | Cat# P10144 |
| BSA | GoldBiotechnology | Cat# A-420 |
| Critical Commercial Assays | | |
| BCA Protein Assay Kit | Pierce | Cat# 23235 |
| Mouse IFN Beta ELISA Kit | pbl | Cat# 42400-1 |
| Human IFN Beta ELISA Kit | pbl | Cat# 41410-1 |
| SsoAdvanced Universal SYBR® Green | Bio-Rad | Cat #1725274 |
| QIAprep spin Miniprep Kit | Qiagen | Cat# 27106 |
| Reasy Mini Kit | Qiagen | Cat# 74104 |
| CellTiter-Blue® Cell Viability Assay | Promega | Cat# G8081 |
| AnnexinV-FITC/PI apoptosis detection kit | BiLegend | Cat# 640914 |
| Clarity Max Western ECL Substrate | Bio-Rad | Cat# 1705061 |
| Exonuclease activity assay kit | VioVision | Cat# K175-100 |
| NE-PER™ Nuclear and Cytoplasmic Extraction Reagents | ThermoFisher Scientific | Cat# 78835 |
| cGAMP ELISA Kit | Cayman chemical | Cat# 501700 |
| Experimental Models: Cell Lines | | |

| | | |
|---|---|---|
| A549 | ATCC | N/A |
| 4T1 | ATCC | N/A |
| CT26 | ATCC | N/A |
| H1299 | ATCC | N/A |
| Experimental Models: Organisms/Strains | | |
| BALB/c | Envigo | Cat # 4702F |
| NOD/SCID | Envigo | Cat # 1700M |
| Software and Algorithms | | |
| GraphPad Prism 8.0 | GraphPad Software, Inc. | https://graphpad.com/scientific-software/prism/ |
| Excel 2016 | Microsoft | https://www.office.com/ |
| ImageJ 1.52a | Wayne Rasband, NIH | https://imagej.net/ |
| BD FACS DIVA 6.2 | BD Biosciences | https://www.bdbiosciences.com/en-us/instruments/research-instruments/research-cell-sorters/facsaria-iii |
| Kaluza | Beckman Coulter Life Sciences | https://www.beckman.com/flow-cytometry/software/kaluza |
| NIS-elements AR 5.02.01 | Nikon | https://www.microscope.healthcare.nikon.com/ |
| Leica TCS SP8 X confocal | Leica | https://www.leica-microsystems.com |

#### Figure S1

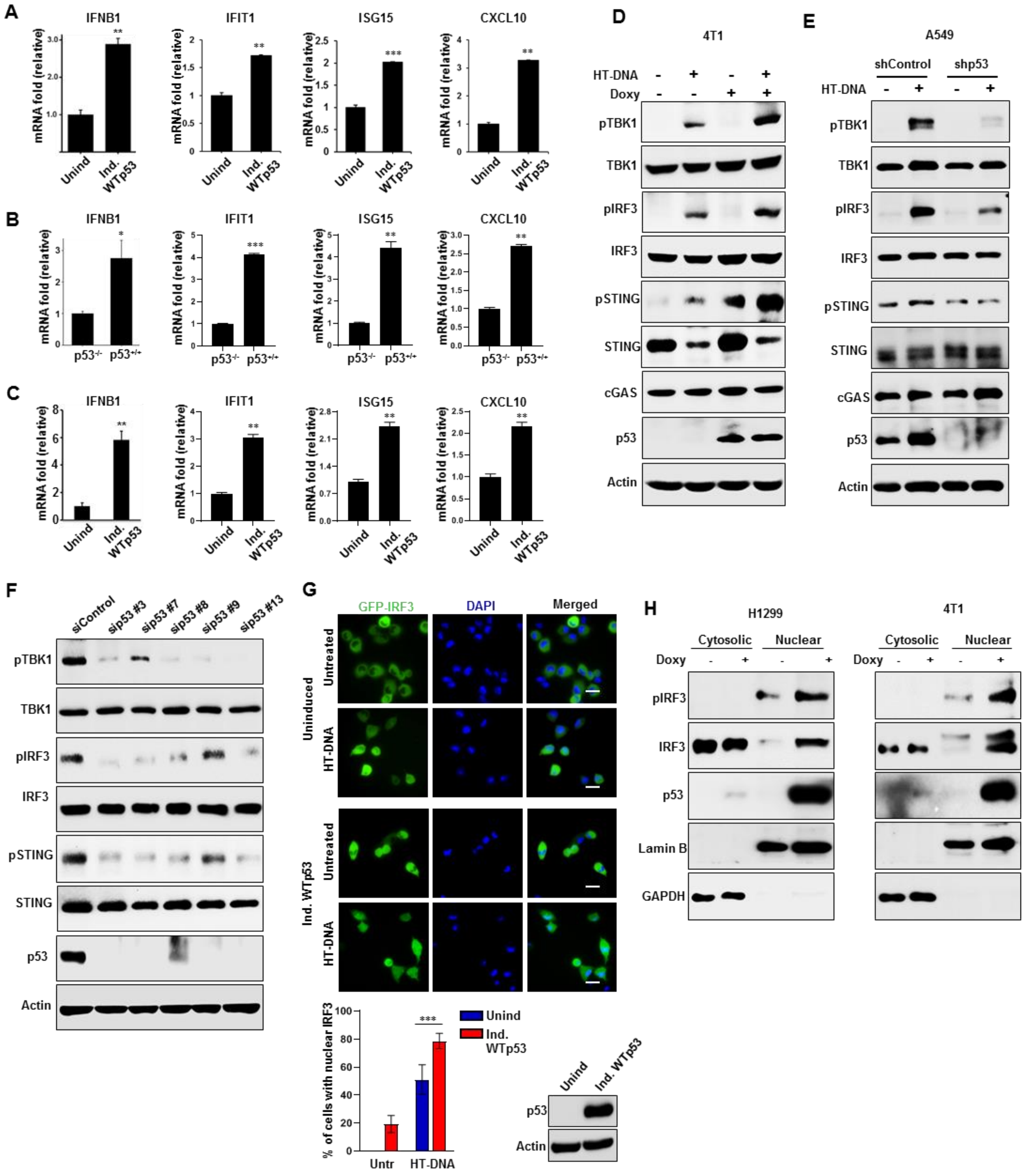
A
IFNB1
IFIT1
ISG15
CXCL10
mRNA fold (relative)
Unind
Ind. WTp53
B
p53-/-
p53+/+
C
D
4T1
HT-DNA
Doxy
pTBK1
TBK1
pIRF3
IRF3
pSTING
STING
cGAS
p53
Actin
E
A549
shControl
shp53
F
siControl
sip53 #3
sip53 #7
sip53 #8
sip53 #9
sip53 #13
G
GFP-IRF3
DAPI
Merged
Uninduced
Ind. WTp53
Untreated
HT-DNA
% of cells with nuclear IRF3
Untr
Unind
Ind. WTp53
H
H1299
Cytosolic
Nuclear
Lamin B
GAPDH

**Figure S1.**

(A-C) Graphs show mRNA expression of IFNB1, IFIT1, ISG15 and CXCL10 in (A) H1299 Induced WTp53 (B) $p53^{+/+}$ and $p53^{-/-}$ MEFs and (C) 4T1 Induced WTp53 cells. (D-E) 4T1 induced WTp53 cells were treated with 2 μg/ml of HT-DNA for 3 h and subjected to western blot analysis. (E) shRNA mediated p53 knockdown (shControl or shp53) A549 cells were treated with 2 μg/ml of HT-DNA for 3 h and cells were harvested for western blotting. (F) A549 cells were transfected with different p53 siRNAs (sip53) and subjected to Immunoblot analysis. (G) Stably GFP-IRF3 positive H1299 cells were engineered to inducibly express WTp53 upon doxycycline treatment. Cells were treated with Doxycycline for 24 h and then treated wit HT-DNA (2 μg/ml) for another 3 h. Representative fluorescence microscopic images are showing IRF3 nuclear localization upon treatment. Scale bar 20 μm. Representative graph shows quantitation of cells having nuclear IRF3 upon HT-DNA treatment. Western blot shows WTp53 induction upon doxycycline treatment. (H) Representative immunoblots of cytosolic and nuclear fractionated lysates of Doxycycline induced WTp53 in H1299 and 4T1 cells. GAPDH and Lamin B were used as loading controls for the cytoplasmic and nuclear fractions, respectively.

Quantification graphs: FoV= Field of View, (n=20) In all panels, error bars represent Mean +/- SD. p values are based on Student's t test. ***$p < 0.001$, **$p < 0.01$.

#### Figure S2

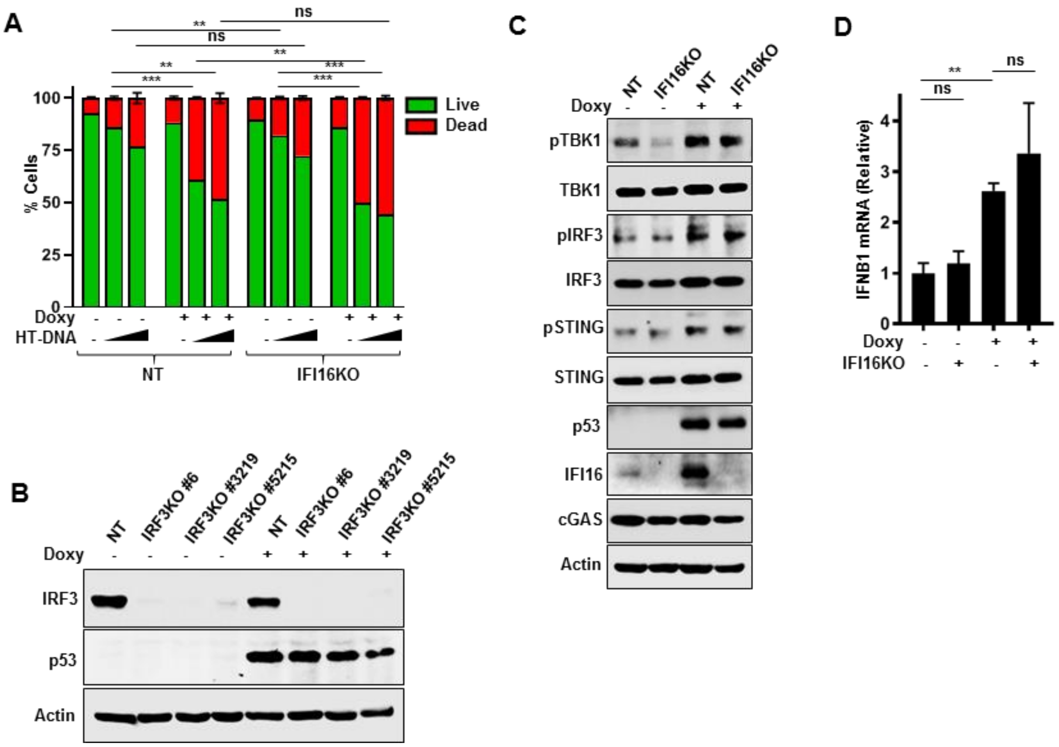


**Figure S2.**

(A) Non-target (NT) and IFI16KO H1299 cells induced WTp53 treated with 2 µg/ml or 4 µg/ml of HT-DNA for 24 h. Cells were harvested, stained with Annexin V-FITC and PI and subjected to flow cytometry analysis. (B) H1299 inducible WTp53 cells were stably knock out for IRF3. Cells were treated with doxycycline to induce WTp53 and subjected to Western blot. (C-D) H1299 inducible WTp53 cells were stably knock out for IFI16. Cells were treated with doxycycline to induce WTp53 and subjected to (C) Western blot or (D) cells were harvested RT-PCR analysis for IFNB1.

Quantification graphs: In all panels, error bars represent Mean +/- SD. p values are based on Student's t test. $***p < 0.001$, $**p < 0.01$, $*p < 0.05$.

#### Figure S3

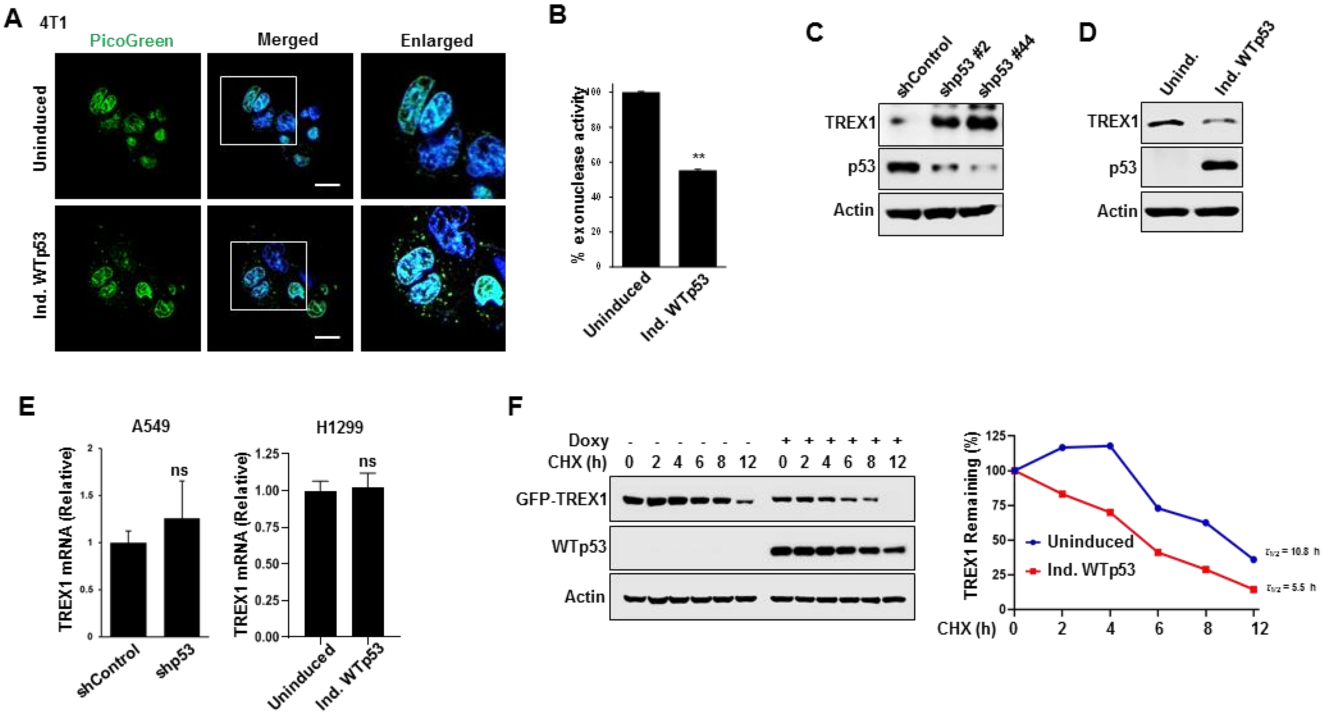


#### Figure S3.

(A) Representative live confocal microscopy images of 4T1 induced WTp53 cells stained for cytosolic DNA using PicoGreen dye and the nucleus was stained with Hoechst 33342. Scale Bar 10 μm. (B) Representative graph indicates % exonuclease activity in 4T1 induced WTp53 cells. (C) CT26 shControl or shp53 cells were subjected to western blot. (D) Representative Immunoblots of 4T1 cells induce WTp53. (E) Representative graphs shows TREX1 mRNA in A549 shControl or shp53 and H1299 induced WTp53 cells. (F) Representative Immunoblots showing H1299 cells were induced to express WTp53 and treated with cycloheximide (20 μM). Cells were harvested at the indicated different time point and subjected to western blot analysis. Representative graph indicate quantification of the relative levels of remaining TREX1 protein after the treatment described in.

Quantification graphs: In all panels, error bars represent Mean +/- SD. p values are based on Student's t test. ***p < 0.001, **p < 0.01, *p < 0.05.

**Figure S4**

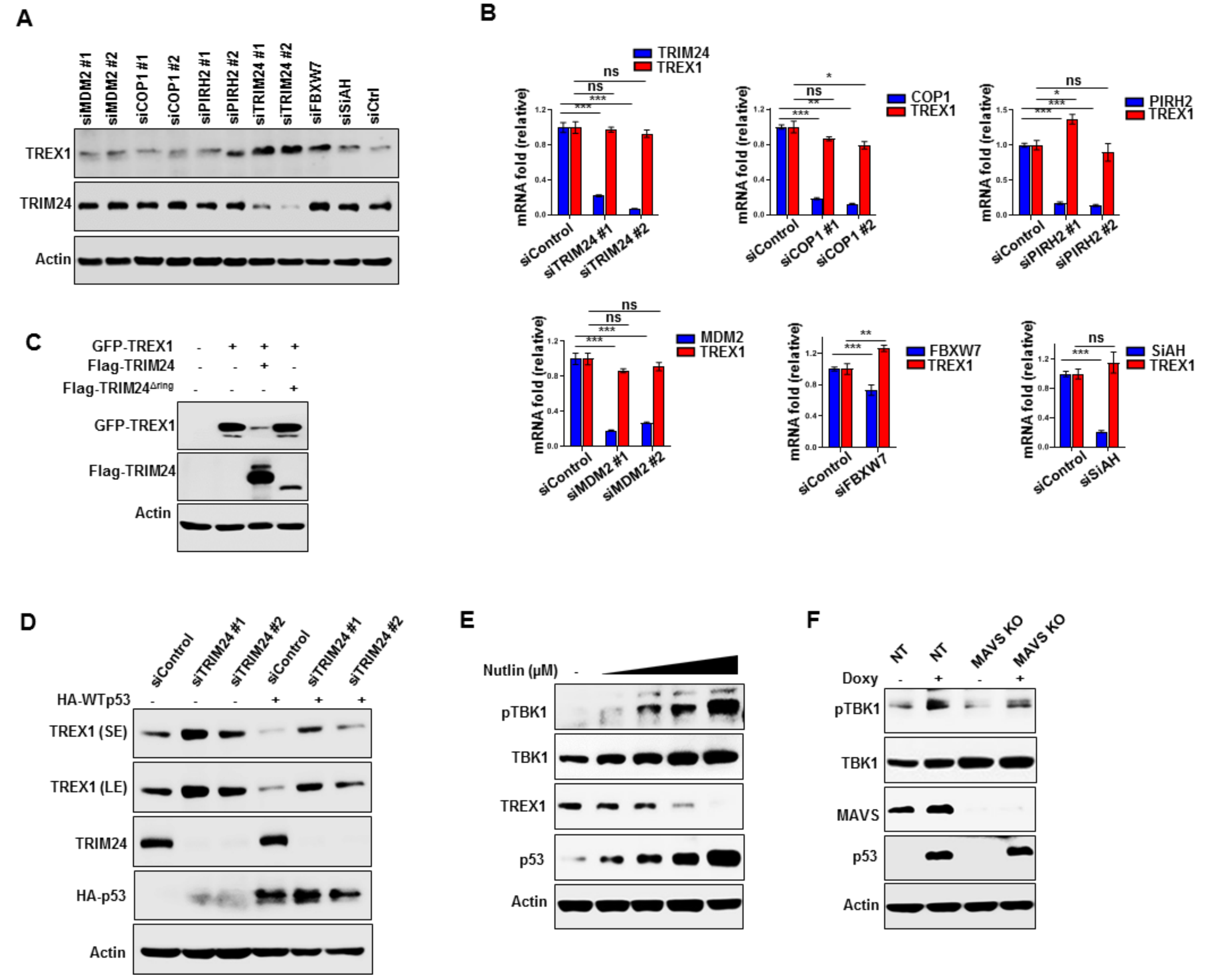


**Figure S4.**
H1299 cells were transfected with siRNAs of MDM2, COP1, PIRH2, TRIM24, FBXW7 and SiAH. Cells were harvested and subjected to (A) western blot analysis or (B) RT-PCR for the indicated genes. (C) H1299 cells were co-transfected with GFP-TREX1 and Flag-TRIM24 or Flag-TRIM24$^{\Delta ring}$ and subjected to western blot. (D) TRIM24 knockdown (siTRIM24) H1299 cells were transfected with HA-WTp53 and subjected to western blot. (E) A549 cells were treated with increasing amount of Nutlin for 24 hrs and cells are subjected to western blot. (F) H1299 inducible WTp53 cells were engineered to CRISPR knockout MAVS and subjected to western blot. Quantification graphs: In all panels, error bars represent Mean +/- SD. p values are based on Student's t test. $^{***}p < 0.001$, $^{**}p < 0.01$, $*p < 0.05$.

### Figure S5

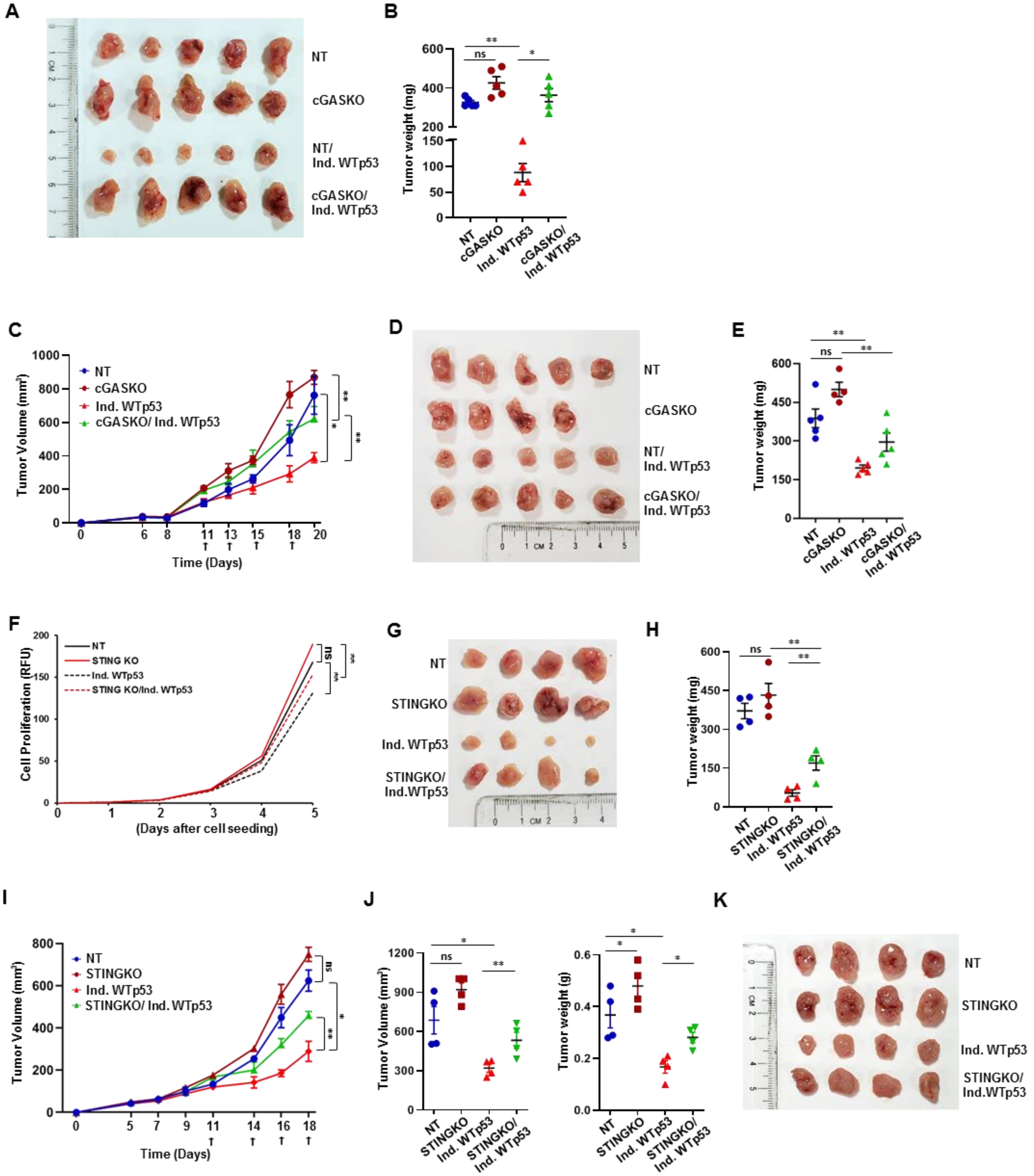

A
NT
cGASKO
NT/ Ind. WTp53
cGASKO/ Ind. WTp53
B
Tumor weight (mg)
ns
**
*
NT
cGASKO
Ind. WTp53
cGASKO/ Ind. WTp53
C
Tumor Volume (mm³)
Time (Days)
NT
cGASKO
Ind. WTp53
cGASKO/ Ind. WTp53
D
NT
cGASKO
NT/ Ind. WTp53
cGASKO/ Ind. WTp53
E
Tumor weight (mg)
F
Cell Proliferation (RFU)
(Days after cell seeding)
NT
STING KO
Ind. WTp53
STING KO/Ind. WTp53
G
NT
STINGKO
Ind. WTp53
STINGKO/ Ind.WTp53
H
Tumor weight (mg)
I
Tumor Volume (mm³)
Time (Days)
NT
STINGKO
Ind. WTp53
STINGKO/ Ind. WTp53
J
Tumor Volume (mm³)
Tumor weight (g)
K
NT
STINGKO
Ind. WTp53
STINGKO/ Ind.WTp53

**Figure S5.**

(A) Representative images showing tumor volume difference of 4T1 induced WTp53 non-target (NT) or cGAS knockout cells in BALB/c mice. (B) Graphical quantification of tumor weight and volume on day 23 in the indicated 4T1 tumor cohorts in BALB/c mice (n= 4). (C) 5 x $10^4$ 4T1 induced WTp53 non-target (NT) or cGAS knockout cells were injected into the of mammary gland of immunodeficient NOD/SCID mice (n=5). Mice were orally given doxycycline to induce WTp53 (indicated by arrow). All mice were sacked on day 21 and graphical quantification represents the tumor growth rate. (D) Representative images showing tumor volume difference of 4T1 induced WTp53 non-target (NT) or cGAS knockout cells in NOD/SCID mice. (E) Graphical quantification of tumor weight and volume on day 21 in the indicated 4T1 tumor cohorts in NOD/SCID mice (n= 5). (F) Representative graphical quantification of *in vitro* cellular proliferation rate between indicated 4T1 cells. (G) Representative images showing tumor volume difference of 4T1 induced WTp53 non-target (NT) or STING knockout cells in immunocompetent BALB/c mice (n=5). (H) Graphical quantification of tumor weight on day 23 in the indicated 4T1 tumor cohorts in BALB/c mice (n= 4). (I) 5 x $10^4$ 4T1 induced WTp53 non-target (NT) or STING knockout cells were injected into the of mammary gland of immunodeficient NOD/SCID mice (n=4). Mice were orally given doxycycline to induce WTp53 (indicated by arrow). All mice were sacked on day 21 and graphical quantification represents the tumor growth rate. (J) Graphical quantification of tumor volume and weight on day 19 in the indicated 4T1 tumor cohorts in NOD/SCID mice (n= 4). (K) Representative images showing tumor volume difference of 4T1 induced WTp53 non-target (NT) or STING knockout cells in immunodeficient NOD/SCID mice (n=4).

Quantification graphs: In all panels, error bars represent Mean +/- SEM. p values are based on Student's t test. $^{***}p < 0.001$, $^{**}p < 0.01$, $*p < 0.05$.

### Figure S6

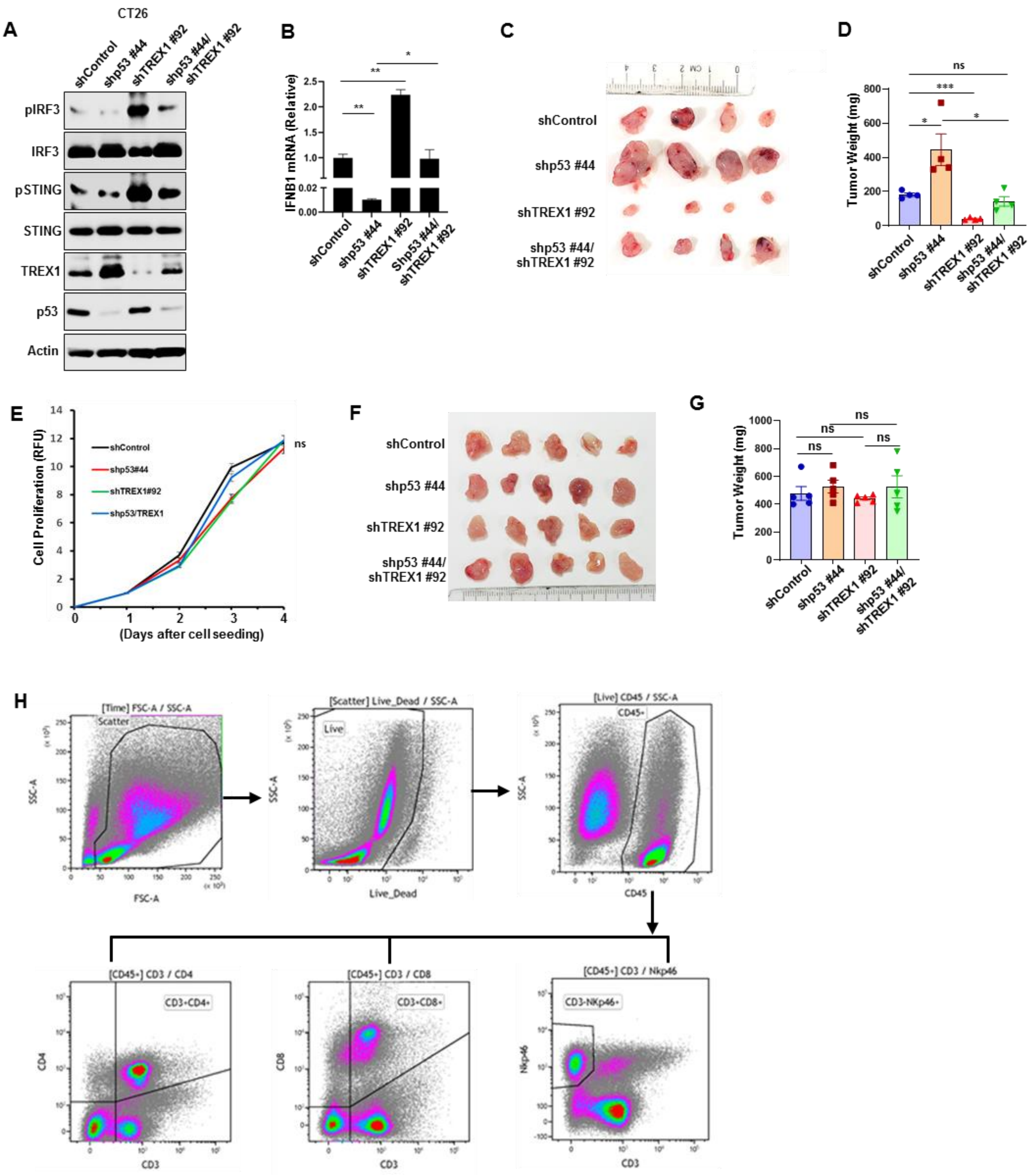
A
CT26
shControl
shp53 #44
shTREX1 #92
shp53 #44/ shTREX1 #92
pIRF3
IRF3
pSTING
STING
TREX1
p53
Actin
B
IFNB1 mRNA (Relative)
shControl
shp53 #44
shTREX1 #92
Shp53 #44/ shTREX1 #92
C
shControl
shp53 #44
shTREX1 #92
shp53 #44/ shTREX1 #92
D
Tumor Weight (mg)
ns
***
*
*
E
Cell Proliferation (RFU)
shControl
shp53#44
shTREX1#92
shp53/TREX1
ns
(Days after cell seeding)
F
shControl
shp53 #44
shTREX1 #92
shp53 #44/ shTREX1 #92
G
Tumor Weight (mg)
ns
H
[Time] FSC-A / SSC-A
Scatter
SSC-A
FSC-A
[Scatter] Live_Dead / SSC-A
Live
Live_Dead
[Live] CD45 / SSC-A
CD45+
CD45
[CD45+] CD3 / CD4
CD3+CD4+
CD4
CD3
[CD45+] CD3 / CD8
CD3+CD8+
CD8
[CD45+] CD3 / Nkp46
CD3-NKp46+
Nkp46

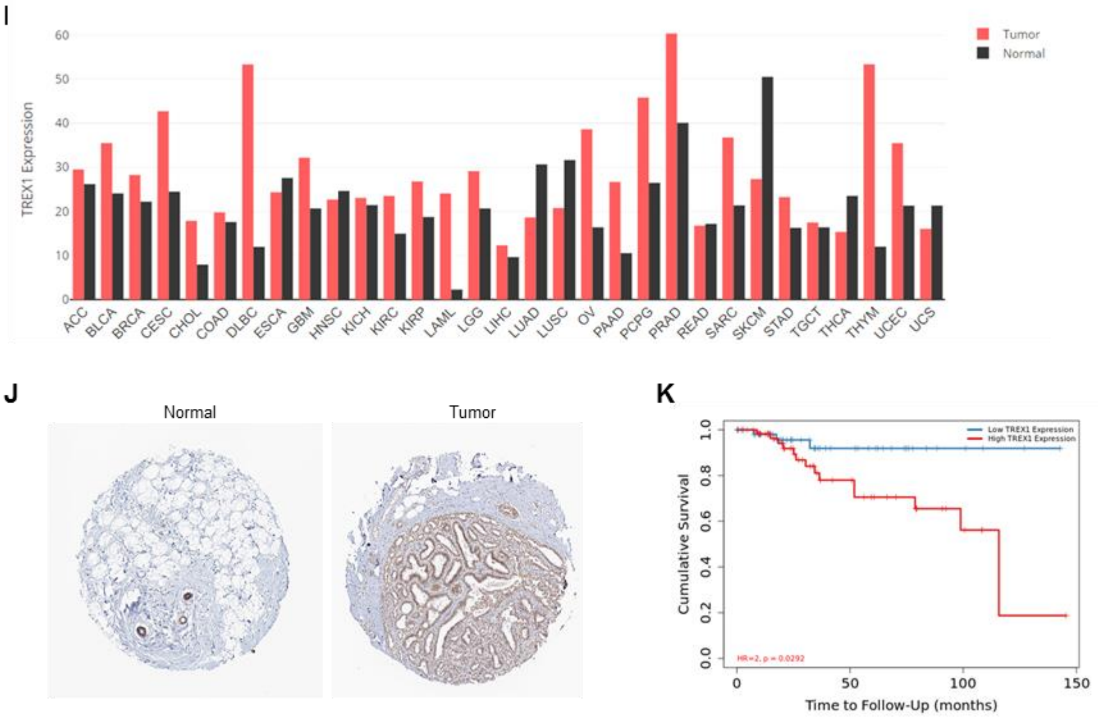


**Figure S6.**

(A) CT26 cells were infected to knockdown either p53 or TREX1 alone or together and subjected to western blot analysis. (B) CT26 shp53, shTREX1 or shp53/shTREX1 cells were subjected to RT-PCR analysis. (C) Representative image showing tumor volume difference of CT26 shp53, shTREX1 or shp53/shTREX1 cells in BALB/c mice. (D) Graphical quantification of difference in tumor weight on day 20 in shp53, shTREX1 or shp53/shTREX1 cohorts (n= 4). (E) Representative graphical quantification of *in vitro* cellular proliferation rate between indicated CT26 shp53, shTREX1 or shp53/shTREX1 cells. (F) Representative image showing tumor volume difference of indicated CT26 cohorts in NOD/SCID mice. (G) Graphical quantification of tumor weight on day 14 in CT26 shp53, shTREX1 or shp53/shTREX1 cohorts in NOD/SCID mice (n= 5). (H) Gating strategy to identify lymphoid immune populations. Scatter plots depicting population of live population of cells isolated from tumors based on FSC and SSC. Following the scatter gate, cells were further gated to identify live CD45+ cells. CD45+ cells were plotted on a scatter dot plot to identify CD3+CD4+ population (T-helper), CD3+CD8+ cells (T-cytotoxic) and CD3-Nkp46+ cells (NK). (I) Expression of TREX1 in different tumor types was determined using the GEPIA (http://gepia.cancer-pku.cn/index.html) databases. (J) Immunohistochemical staining of TREX1 protein in Breast cancer tissues and corresponding normal tissues was obtained from the Human Protein Atlas (HPA) database. (K) Kaplan–Meier survival curves comparing the high and low expression of TREX1 in breast carcinomas (BRCA) as determined using the TIMER 2.0 database.

Quantification graphs: In all panels, error bars represent Mean +/- SEM. p values are based on Student's t test. $^{***}p < 0.001$, $^{**}p < 0.01$, $^{*}p < 0.05$.
